# Microbiome restructuring: dominant coral bacterium *Endozoicomonas* species display differential adaptive capabilities to environmental changes

**DOI:** 10.1101/2021.10.31.466697

**Authors:** Kshitij Tandon, Yu-Jing Chiou, Sheng-Ping Yu, Hernyi Justin Hsieh, Chih-Ying Lu, Ming-Tsung Hsu, Pei-Wen Chiang, Hsing-Ju Chen, Naohisa Wada, Sen-Lin Tang

## Abstract

Bacteria in the coral microbiome play a crucial role in determining coral health and fitness, and the coral host often restructures its microbiome composition in response to external factors. An important but often neglected factor determining this microbiome restructuring is the capacity of microbiome members to adapt to a new environment. To address this issue, we examined how the microbiome structure of *Acropora muricata* corals changed over 9 months following a reciprocal transplant experiment. Using a combination of metabarcoding, genomics, and comparative genomics approaches, we found that coral colonies separated by a small distance harbored different dominant *Endozoicomonas related phylotypes belonging to two different* species, including a novel species, *Candidatus* Endozoicomonas penghunesis 4G, whose chromosome level (complete) genome was also sequenced in this study. Furthermore, the two dominant *Endozoicomonas* species showed varied adaptation capabilities when coral colonies were transplanted in a new environment. The differential adaptation capabilities of dominant members of the microbiome can a) provide distinct advantages to coral hosts when subjected to changing environmental conditions and b) have positive implications for future reefs.

## Introduction

Bacteria are one of the main microbial partners in the coral holobiont [1]. They may play a role in coral health, disease, and nutrient supply [2, 3]. A coral colony often accommodates several hundred, if not thousands, of bacterial phylotypes [4, 5], with different bacterial communities residing in coral compartments such as the coral mucus [6–8], tissue [6, 7, 9, 10], gastrovascular cavity [11], and skeleton [12–14]. These bacterial communities are often diverse, dynamic, and—according to many studies—profoundly influenced by factors such as host specificity and spatiotemporal changes in the surrounding environment [9, 15–19].

Of the factors involved in restructuring the coral-associated bacterial community, environmental changes and host specificity are two major drivers influencing the composition of the bacterial community in corals. In terms of environmental changes, numerous studies have reported shifts in the bacterial community composition of corals in response to variations in temperature [20–24], nutrient load [21, 25], exposure to pathogens [26], and anthropogenic factors [21, 27]. Regarding host specificity, the same coral species living in habitats hundreds to thousands of kilometers apart were found to accommodate similar bacterial community profiles [1, 28], whereas adjacent corals colonies of different species had distinct microbiomes [1, 29]. Interestingly, several studies have asserted that changes to the coral microbiome composition in response to new environments are host-specific; this was tested via transplantation experiments and suggests that microbiome alteration is a potential acclimatization strategy [22, 30]. This microbiome alteration potential is known to vary depending on the host species. For example, Ziegler and coworkers [30] studied variation in the microbiomes of the corals *Acropora hemprichii* and *Pocillopora verrucosa* in a long-term cross-transplantation experiment and identified that *A. hemprichii* harbors a highly flexible microbiome, whereas *P. verrucosa* has a remarkably stable microbiome, even after exposure to different levels of chronic pollution, suggesting that the bacterial communities of different coral species vary.

In most of the studies conducted to date, factors influencing the changes in the coral-associated bacterial community are often external, such as those mentioned above. Only recently have internal factors like host genotype been shown to also influence the coral microbiome [31]. However, the adaptation capability of bacteria, one hidden but crucial internal factor, has long been neglected. Theoretically, based on the nature of genetic variations among bacteria, some bacterial phylotypes of the same bacterial group in a community may perform better than others under specific environmental conditions due to higher adaptation capabilities. Therefore, those bacteria phylotypes with higher adaptation capacities could maintain a more stable abundance profile during specific environmental changes and potentially play important functional roles. In other words, along with host selection and environmental influence, changes in the bacterial community may be greatly affected by the adaptation capacities of individual bacterial groups. However, this aspect of microbiome restructuring is mostly unexplored.

To test the above hypothesis that bacterial groups have different adaptation capabilities, we first needed to identify a dominant bacterial group often identified in corals with multiple operational taxonomic units (OTUs) or, more recently, amplicon sequence variants (ASVs) from metabarcoding surveys. One such group belongs to the genus *Endozoicomonas* (Phylum: *Proteobacteria*, Class: *Gammaproteobacteria*, Order: *Oceanospiralles*, family: *Endozoicomonadecea* (also *Hahellacea*)), a dominant bacterial group found in several coral species [32, 33]; it plays a role in coral health and nutrition regulation [34]. In a recent study from our group, Shiu et al. [23] that the abundance profiles of certain *Endozoicomonas* OTU shifted within 12 hours under thermal stress. If *Endozoicomonas* does have a differential adaptation capability, then we can hypothesize that different *Endozoicomonas* phylotypes behave differently and some may remain more stable and colonize longer than others when corals are subjected to environmental change. Another prerequisite to testing the differential capability of bacterial phylotypes is finding a region with differential environmental conditions within a small distance such that geographical variation does not influence the coral microbiome.

The Penghu Archipelago, located in the Taiwan Strait, has been proposed to be a climate-change refugium for corals and has a unique thermal regime, governed by the warm Kuroshio Current in the summer and cold China Coastal Current in the winter [23, 35]. These factors make it an ideal location for an experimental site. The semi-closed Chinwan Inner Bay (hereafter “Inner Bay”) of Penghu has suffered substantial marine biodiversity losses, including significant damage to marine aquaculture, wild fisheries, and coral bleaching due to extreme weather events in the winter [36]. Furthermore, based on regional news and government reports, domestic sewage dumping, the presence of a shipping port, and aquaculture practices have increased the concentrations of nitrogen and ammonia in the calmer waters of the Inner Bay compared to the Outer Bay region, threatening corals. The contrasting local environmental conditions between the Inner and Outer Bays make them excellent sites to study the response of locally acclimated coral microbial communities (especially for *Endozoicomonas*), trace coral microbiome restructuring at a fine scale, and test the differential adaptability hypothesis of dominant coral-associated bacteria.

We examined how the microbial community restructures in response to changes in the local environment and tested the hypothesis that *Endozoicomonas* phylotypes have a differential adaptation capacity using colonies of the coral *Acropora muricata* (genus: *Acropora*) in the Penghu Archipelago, Taiwan. These coral species in the Penghu Archipelago have been reported to harbor *Endozoicomonas* as their dominant bacteria [23]. We conducted a longitudinal (9-month) *in situ* reciprocal transplant experiment with repeated sampling, where coral colonies from the semi-closed Inner Bay were transplanted into the open ocean region of the Outer Bay and vice versa. Furthermore, we aimed to isolate, culture, and characterize dominant *Endozoicomonas* members to provide genomic insights into how bacteria adapt to these environments.

## Materials and methods

### Study design and experimental setup

Five colonies of *Acropora muricata* (40×40 cm) were collected at a depth of 3 m from the Outer Bay (O) (N23° 33.097’ E119° 38.335’) and Inner Bay (I) (N23° 31.853’ E119° 33.629’) along the reef adjacent to the coast of the Penghu Archipelago, Taiwan (Figure 1A). Mother coral colonies were first collected in April. These acted as controls for the native coral microbiome in the study sites (Figure 1A). Later, mother colonies were fragmented into two halves (approx. 20×20 cm each). Coral fragments from mother colonies were either cross swapped (I→O or O→I) or transplanted in their original location (IC or OC) (Figure 1B). Coral fragments from each colony that remained in their original location acted as controls to measure any change in the microbiome due to the transplant procedure and change in the microbiome based on colony age and experimental time. Coral fragments were glued onto the reef with epoxy putty.

**Figure 1.**
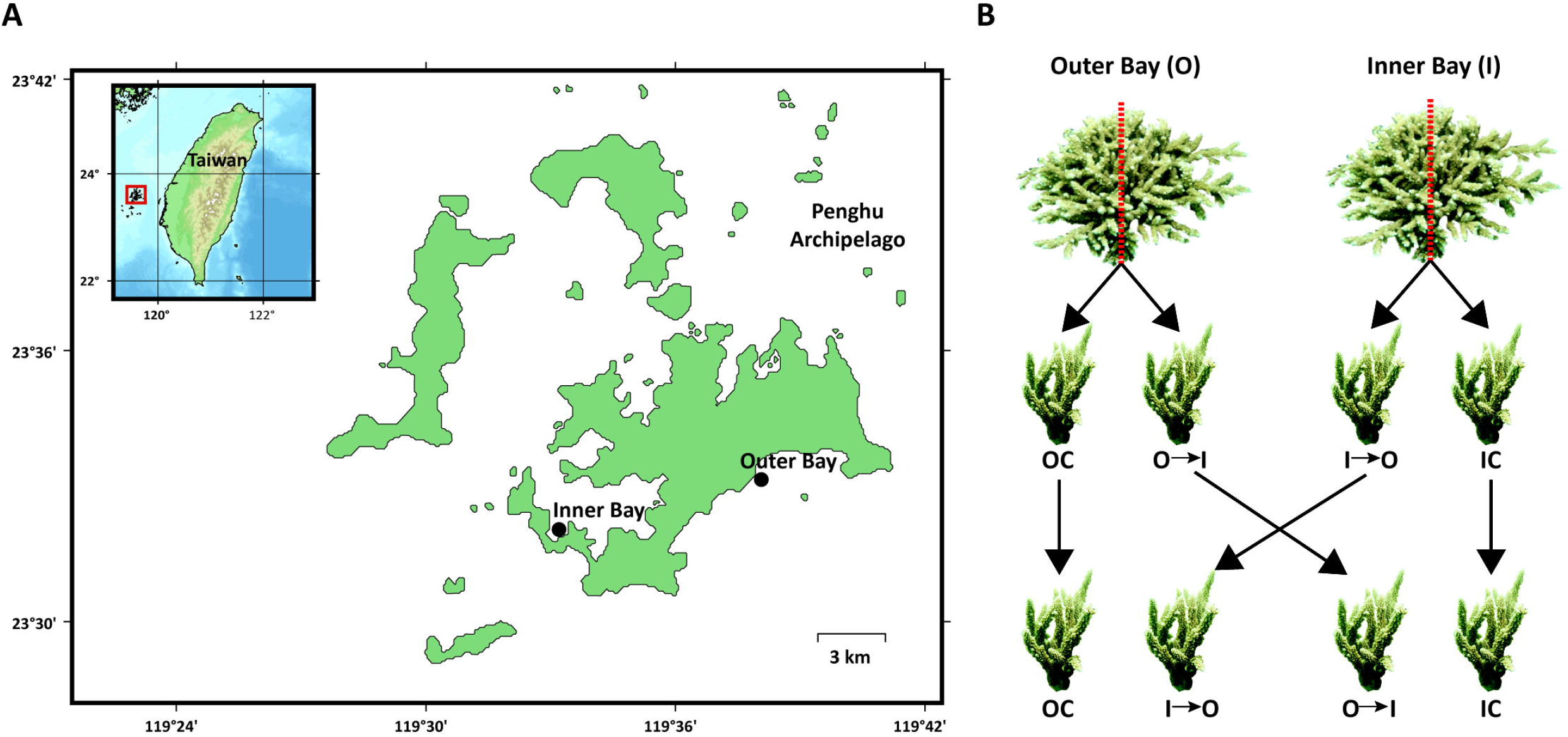
Sampling location and reciprocal transplant experiment overview. **A)** Map of the Penghu Archipelago, Taiwan with two sampling sites: Inner Bay and Outer Bay. **B)** Schematic representation of the reciprocal transplant experiment setup with sample codes OC: Outer Bay Control; IC: Inner Bay Control, O→I: Outer Bay colonies transplanted into the Inner Bay and I→O: Inner Bay colonies transplanted into the Outer Bay.

### Sampling timeline and sample collection

Study sites were visited every month from April to August 2018 and then in December 2018 to check the status of transplanted fragments and collect samples. In total, we collected 122 samples, including seawater samples (1 L) at each time-point and location. 2×2 cm fragments were taken from each colony, rinsed with filtered seawater, and stored in 99% ethanol at -20°C before DNA extraction.

### DNA extraction and 16S rRNA gene amplicon sequencing

Frozen coral fragments were sprayed (70 psi) with ∼15 ml 1x TE buffer (10 mM Tris-HCl, 1 mM EDTA, pH 8), then placed into sterile zip lock bags. Total DNA was extracted using a modified CTAB method [37]. Coral tissue samples DNA was extracted with conventional Chloroform/isoamyl alcohol (24:1) and phenol/chloroform/isoamyl alcohol (25:24:1) step and isopropanol precipitation method. The DNA pellet was rinsed with 70% ethanol and then dissolved in 50 µl ddH_2_O and stored at -20°C. DNA concentration was determined using a NanoDrop 1000 Spectrophotometer (Thermo Fisher Scientific, Waltham, MA, USA) and Quant-iT dsDNA HS (High-Sensitivity) Assay Kit. Seawater samples were processed similarly with the modified CTAB method [37].

For DNA library construction, 968F (5’-AAC GCG AAG AAC CTT AC-3’) [38] and 1391R (5’-ACG GGC GGT GWG TRC-3’) [39] universal primers were used to amplify the bacterial V6-V8 hypervariable region of the 16S rRNA gene from the total DNA from samples using PCR. For PCR reactions, 50–150 ng of template DNA were used. PCR was performed in 50 µl reaction volumes, consisting of 1.5 U *TaKaRa Ex Taq* (Takara Bio, Otsu, Japan), 1X *TaKaRa Ex Taq* buffer, 0.2 mM deoxynucleotide triphosphate mixture (dNTPs), 0.2 mM forward and reverse primers, and template DNA. The PCR conditions consisted of an initial denaturing step at 95°C for 5 min; followed by 30 cycles at 94°C for the 30 s, 52°C for 20 s and 72°C for 45 s; and a final extension at 72°C for 10 min. The amplified product was visually confirmed using 1.5% agarose gel with a 5 µl PCR product. Target bands (∼420 bp) were cut and eluted using a QIAEX II gel extraction kit (Qiagen, Valencia, CA, USA).

Each bacterial V6-V8 amplicon was tagged with a unique barcode sequence by designing tag primers with 4-base overhangs at 5’ ends. The sample-specific tagging reaction was performed with a 5-cycle PCR, with a reaction program of initial denaturation at 94°C for 3 min, followed by denaturation at 94°C for 30 s, annealing at 52°C for 20 s, extension at 72°C for 45 s, and a final extension at 72°C for 10 min. The amplified product was purified using the QIAquick PCR purification Kit (Qiagen, Valencia, CA, USA). Purified products were pooled into four independent libraries and sequenced with Illumina MiSeq paired-end sequencing (2×300) at Yourgene Biosciences, Taiwan.

### Sequence data processing and analysis

Paired-end raw reads obtained from Illumina sequencing were merged using USEARCH v11 [40] with the parameters minovlen=16, maxdiffs=30, and pctid=80. Merged reads were sorted, quality-filtered, and trimmed using Mothur v1.3.81 [41]. Reads 400–470 bp long with an average quality >25 were kept. Chimeric reads were inspected and eliminated with UCHIME [42] by USEARCH v11. Qualified reads were retained for subsequent analysis. High-quality reads were denoised using UNOISE3 [43], and zero-radius Operational taxonomic units (zOTUs)—which are equivalent to exact sequence variants—were obtained. The denoised sequences were aligned against the SILVA128 [44, 45] ribosomal RNA database for a taxonomic assignment up to the genus level using Mothur on a per-sample basis with a pseudo-bootstrap cutoff of 80%.

### Statistical analyses

All statistical analyses and graphs were generated in R (R Core Team 2020). Stacked bar plots were obtained by converting absolute abundance profiles into relative abundances. Abundance profiles were processed with the R packages phyloseq [46], vegan [47], ggplot2 [48], pheatmap [49], and microbiomeMarker [50] for downstream analyses and visualization. Alpha diversity analysis was conducted after rarifying the samples to an even depth of 5,704 reads using the estimate_richness function from phyloseq. Alpha diversity metrics were compared using Analysis of Variance (ANOVA) and Tukey’s post hoc tests using vegan package *p-*value correction for multiple testing. Multivariate analysis was performed after square-root transforming the zOTUs count data. Betadisper function was used to calculate the multivariate dispersion of samples (Bray-Curtis distance) between sample groups. Homogeneity of multivariate dispersion was tested with ANOVA. Non-multidimensional scaling (nMDS) was performed to compare community compositions using the Bray-Curtis distance metric between sample groups. Permutational multivariate analysis of variance (PERMANOVA) with the “adonis” function (with 9,999 permutations) was used to statistically test for differences in community compositions between the back and cross transplant samples for each location as dispersion was significantly different between groups. Linear discriminant analysis Effect Size (LEfSe) implemented in the microbiomeMarker package in R was used to identify shifts in zOTUs between back and cross transplant samples for each location with a log(LDA) cutoff of 3 (Kruskal-Wallis test: p <0.05). z-score transformed abundance profiles of marker zOTUs identified from LEfSe were visualized with a heatmap via pheatmap.

### Environmental parameters

The water temperatures of the Outer (O) and Inner (I) Bay of the Penghu Archipelago, Taiwan were obtained from May 2018 through to December 2018 using temperature data loggers (HOBO© Pendant, Onset Corp, United States) located at ∼3 m deep, close to target colonies, and recording temperatures every 30 min. Abiotic factors including NH_3_, NO_3,_ and PO_4_ were measured with LaMotte 1910 SMART^®^3 Colorimeter; pH was measured with a HORIBA LAQUA act water quality meter; and salinity was measured with an ATAGO master refractometer.

### Bacteria isolation and culturing

*Ca.* Endozoicomonas penghunesis 4G was isolated from the coral *Acropora muricata* off the coast of the Inner Bay, Penghu Archipelago, Taiwan (GPS location: N23° 31.851’ E119° 33.631’). Coral tissue and mucus were sprayed with TE buffer (10 mM Tris-HCl, 1 mM EDTA, pH 8) and serially diluted to 10^-4^. All dilutions were plated on Modified Marine Broth version 4 (MMBv4 agar) (Ding et al. 2016) and incubated at 25°C. Each colony was screened first by the following primers: bacterial universal forward 27F (5’-AGA GTT TGA TCM TGG CTC AG-3’) and *Endozoicomonas*-specific reverse En771R (5’-TCA GTG TCA RRC CTG AGT GT-3’) [51]. *Endozoicomonas* sp. 16S rRNA gene V1-V4 region was PCR amplified by 35 cycles with a denaturing step at 94°C for the 30 s, followed by annealing at 54°C for 30 s and an extension step at 72°C for 45 s. PCR product was checked on a 1.5% agarose gel after electrophoresis. All samples with bands ∼750 bp long were then sub-cultured in MMB medium. Full-length 16S rRNA genes were amplified by universal bacterial primer 27F (5’ – AGA GTT TGA TCC TGG CTC AG- 3’) and 1541R (5’- AAG GAG GTG ATC CAG CC -3’). The full-length 16S rRNA PCR reaction was amplified using 30 cycles with a denaturing step at 95°C for 30 s, annealing at 55°C for 30 s, and a final extension at 72°C for 90 s. Amplified products with target bands (∼1465 bp) were cut and later sequenced by Sanger sequencing (3730 DNA analyzer, Thermo, USA) from Genomics, Taipei, Taiwan. Chromatograms obtained were manually checked and sequences were trimmed. The final length of the high-quality trimmed sequence was ∼600 bp. Sequences with ≤98% identity to 16S rRNA genes of type strains from genus *Endozoicomonas* were deemed new candidates for novel *Endozoicomonas* species.

### Physiological characterization

*Ca.* E. penghunesis 4G was cultivated on a Modified Marine broth version 4 (MMBv4 medium) [52] (Table S1) for enrichment and a broad range of physiological characterizations was performed. The optimum salinity was tested on MMB medium with NaCl concentrations adjusted as required (0.5% and 1.0 ∼ 4.0%, w/v, in increments of 1.0%). The growth temperature range was tested at 4°C and 10–40°C (at 5°C intervals). The pH tolerance was determined using the following buffers: pH 4.0–7.0, HCl; pH 7.0–10.0, and NaOH (at 1.0 pH unit intervals).

Three physiological tests (pH, salinity, and temperature) were measured based on the turbidity (at OD_600_) of cultures grown at different pH values, NaCl concentration, and temperatures, respectively. Commercial API 20NE kits (bioMérieux, France) were used to test the ability to metabolize different carbon substrates per the manufacturer’s protocol. Additional carbon utilization was evaluated in Modified Marine medium (see details in Table S2). Bacterial motility was tested in Marine Broth semisolid agar (0.5% agarose). The Gram stain kit (Fluka, England) was used to distinguish bacterial Gram reactions. Relation to oxygen was determined after incubating *Ca.* E. penghunesis 4G on MMB agar in the 2.5L Oxoid AnaroGen system (Thermo, USA) and cultured at 25°C for 7 days. Oxidase and catalase activity was tested independently by adding 35% H_2_O_2_ and 0.1% tetramethyl--phenylenediamine dihydrochloride, respectively. An antibiotic sensitivity test was performed after spreading bacteria on an MMB plate with each disc containing different antibiotics (10 µg streptomycin and 10 µg ampicillin). The results were observed after 5 days of incubation at 25°C, and sensitivity was measured based on the distance from the discs to the edge of the clear zone. Bacteria were scored as resistant if the diameter was greater than 2 mm, slightly sensitive if the diameter was 1–2 mm, and resistant otherwise.

### Morphological characterization

The morphology of *Ca.* E. penghunesis 4G, including colony shape and color, was observed by a stereomicroscope (Leica EZ4, Germany). General cell structure and cell Inner structure were studied by transmission electron microscopy (TEM). The bacterial shape on a single colony was observed by scanning electron microscopy (SEM). TEM and SEM observations were made after bacteria were cultured in MMB for 1 day and MMB agar (1.5%) for 3 days, respectively. Colonies were incubated at 25°C.

For the *Ca.* E. penghunesis 4G thin section, bacteria were first centrifuged at 2500 *g* for 5 minutes and bacterial pellets were collected and fixed in 2.5% glutaraldehyde and 4% paraformaldehyde in a 0.1 M sodium phosphate buffer (pH 7.0) at room temperature for 1 h. After three 20-min buffer rinses, the samples were post-fixed in 1% OsO_4_ in the same buffer for 1 hour at room temperature and then rinsed again as above. Samples were dehydrated in an alcohol series, embedded in Spurr’s resin (EMS, USA), and sectioned with a Leica EM UC6 ultramicrotome (Leica, Germany). The ultra-thin sections (70–90 nm) were stained with 5% uranyl acetate in 50% methanol and 0.4% lead citrate in 0.1 N sodium hydroxide. An FEI G2 Tecnai Spirit Twin TEM (FEI, USA) at 80 kilo-volts for viewing and images were captured with a Gatan Orius CCD camera (Gatan, USA).

The colony of *Ca.* E. penghunesis 4G was observed using cryo-SEM (FEI Quanta 200 SEM/Quorum Cryo System PP2000TR FEI). The MMBv4 agar plate containing a single colony of *Ca.* E. penghunesis 4G was sectioned into 1 mm x 1 mm and loaded onto the medium-containing stub, and then frozen with liquid nitrogen slush. The frozen sample was transferred to the sample preparation chamber at -160°C. After 5 min, the temperature was raised to -85°C, and the samples were etched for 20 min. After coating at -130°C, the samples were transferred to the SEM chamber and observed at -160°C and 20 KV.

The general cell morphology was studied by negative staining and observed under TEM. Bacteria were enriched in MMB for 1 day before adding a fixative solution (2.5% glutaraldehyde + 4% paraformaldehyde/0.1M PBS) at 37°C for 10 min. To reduce the background signal of TEM observation, MMB was replaced first by PBS then by sterilized H_2_O twice, and the bacteria were mounted onto grow-discharge carbon-formvar grids. Bacteria were stained by 2% phosphotungstate for 1 s and, finally, the sample was rinsed with sterilized H_2_O twice and viewed under FEI G2 Tecnai Spirit Twin TEM at 80 KV. The images were then captured with a Gatan Orius CCD camera.

### Phylogenetic analysis of *Endozoicomonas* sp. zOTUs

To phylogenetically place the dominant *Endozoicomonas* zOTUs identified and the 16S rRNA gene of *Ca.* E. penghunesis 4G, and to identify their closest neighbor within genus *Endozoicomonas* and its cultured isolates, representative 16S rRNA sequences from type strains (12 total) and one outgroup *Halospina denitrificans* HGD were downloaded from the NCBI taxonomy database (https://www.ncbi.nlm.nih.gov/taxonomy). Sequences were aligned using the RNA homology search tool cmalign [53] from the infernal package, and the CM models for domain bacteria were acquired from the rfam database [54]. A maximum-likelihood phylogeny tree was built using the IQ-TREE web server [55] with 1000 bootstraps and best model selection enabled (best model: K2P+I+G4). The tree was finally visualized and edited in the laboratory-licensed version of iTOL v4 [56].

### Long- and short-read paired-end sequencing and genome assembly

Long reads obtained from nanopore sequencing were first quality checked with nanoqc [57] and only high-quality paired-end reads were used for genome assembly using metaFlye [58] with default settings. Illumina reads (2x 300) were first quality checked with FastQC (https://www.bioinformatics.babraham.ac.uk/projects/fastqc/), then the adapters were removed and the reads trimmed with AdapterRemoval v2 [59]. High quality (phred >30) and trimmed paired-end reads were used to polish the crude nanopore assembly with four rounds of pilon [60] with default settings.

### Genome annotation

The assembled genome was first checked for completeness, contamination, and heterogeneity using CheckM [61]. The *E. acroporae* Acr-14^T^ genome [62] was assembled previously in our laboratory. Protein predictions in two *Endozoicomonas* sp. was performed with Prodigal in Prokka [63] with default settings to keep gene calls preserved for further functional categories analysis. A Rapid Annotation using subsystem technology (RAST) server [64] was used to obtain higher-order subsystem level features. The “reconstruct pathway” approach in blastKOALA v2.2 [65] was used to obtain KEGG Ontology (KO) terms and in-depth annotation of the proteome. CRISPRcasFinder [66] was used to access the CRISPR-spacer. Eukaryote-like proteins (ELPs) were searched from a Batch Web-CD search against the CDD database [67], with minimum e-value 1e-5 and maximum hit number set to 50. Circular genomic map of *Ca.* E. penghunesis 4G was visualized by CGView Server beta [68].

### Data availability

All sequencing data generated in this manuscript was submitted under the BioProject: PRJNA758232 and the *Ca.* E. penghunesis genome was made available under Accession ID: SAMN21016876

## Results

### Sampling and sequencing overview

We collected a total of 110 coral and 12 seawater samples from the experiment, of which 10 coral fragments were removed (all from the Inner Bay) during the experiment (marked with an ‘X’ in Figure 2A–B) as they appeared to be dead. These dead samples were only used to help contrast with the microbial community compositions of healthy corals and were later removed before downstream analyses, including α and β diversity analyses. At the end of the experiment, we had 100 coral and 12 seawater samples. A total of 2,015,935 high-quality reads (average 16,524 reads per sample) were obtained after removing chimeras and poor-quality reads from the 110 coral and 12 seawater samples. These reads were denoised into 2064 zOTUs. Healthy corals (n=100) had 1,815,002 reads (range: 6092–67598), 1966 zOTUs and seawater (n=12) had 149,638 reads (range: 6897–19777) and 1521 zOTUs.

**Figure 2.**
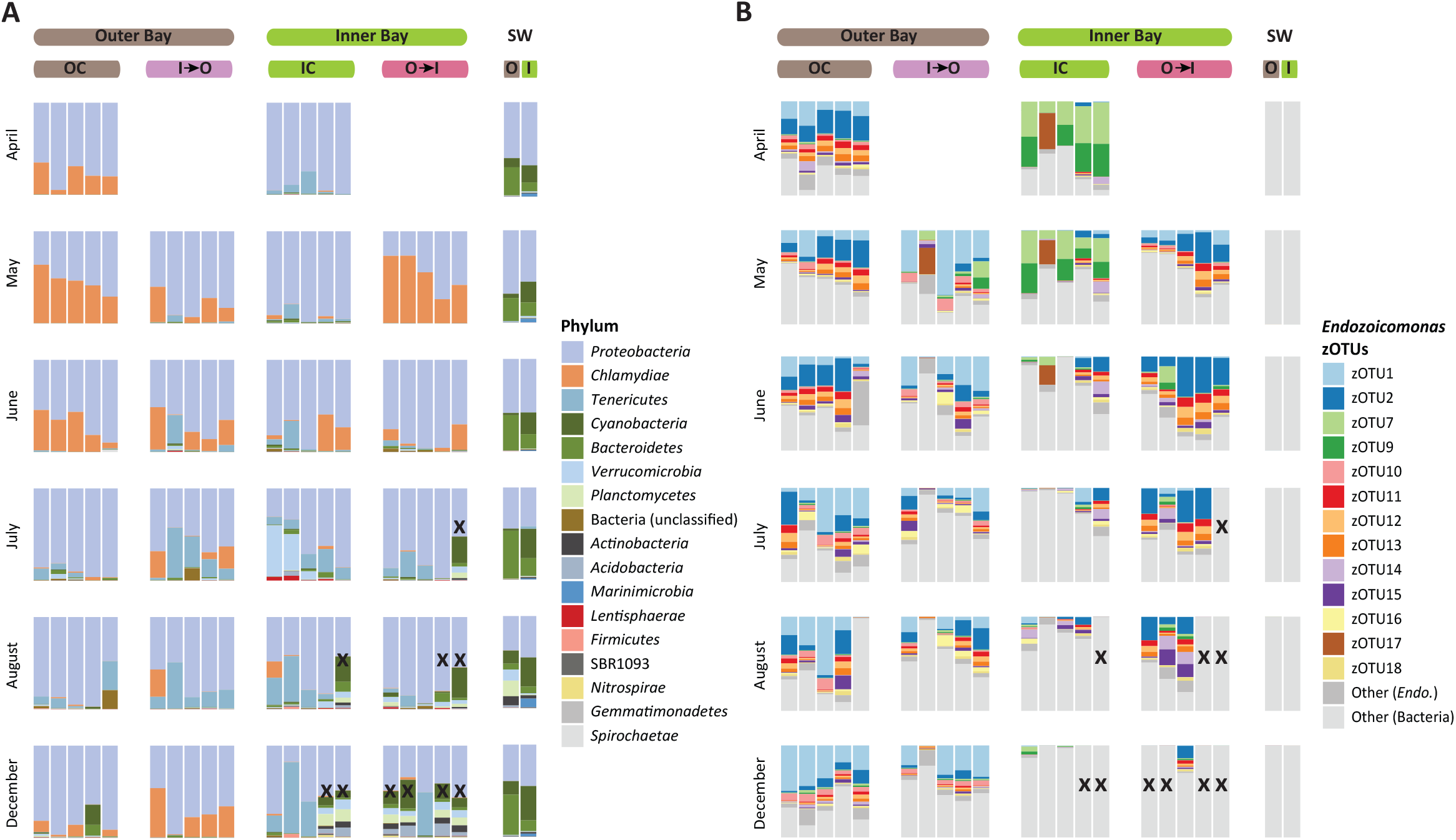
Bacterial community composition overview. **A)** Relative abundance-based bacterial community composition at the phylum level across all sample sets (IC, OC, I→O and O→I). **B)** Relative abundance of different Endozoicomonas zOTUs across all sample sets. “X” in the figure denotes dead colonies.

### Coral and seawater microbiomes differ in bacterial diversity and compositions

The coral and seawater samples were significantly different in bacterial diversity and evenness, measured through zOTU richness (Figure S1A), Shannon (Figure S1B) and Chao1 (Figure S1C) diversities, and Inverse Simpson evenness (Figure S1D). The seawater samples had more than twice the zOTU richness and Shannon and Chao1 diversities compared to the coral samples. August samples showed an unusual alpha diversity pattern, particularly in seawater samples; this could be because the samples were collected during heavy rainfall (±3 days). We observed an increase in richness and Chao1 between O→I and OC samples, but no apparent differences were observed between I→O and IC samples. In terms of corals at different locations, there was no significant difference in calculated alpha diversity measures between control and transplant samples from the Inner and Outer Bay (Figure S1 A–D).

*Proteobacteria* (specifically Class *Alphaproteobacteria*) and *Bacteroidetes* (Class: *Flavobacteriia*) were the dominant phyla in seawater samples across all time points, followed by *Cyanobacteria*, which was particularly abundant in Inner Bay samples. There was heavy rainfall during the week in August that samples were collected, and we noticed a higher abundance of *Marinimicrobia* (Class: Marnimicrobia SAR406 clade), *Planctomycetes,* and *Verrucomicrobia* in those samples, which might explain the different bacterial diversity and evenness results obtained that month. On the contrary, coral samples were dominated by *Proteobacteria* (specifically Class: *Gammaproteobacteria*)*, Chlamydiae,* and *Tenericutes* (Class: *Mollicutes*). Dead coral samples had a similar bacterial community composition as seawater samples (Figure 2A, Figure S2).

### Changes in the coral microbial community throughout the reciprocal transplant

*Proteobacteria* was the dominant phylum across all sample groups (control: IC and OC; transplant: I→O and O→I) from the two locations throughout the experiment. We observed shifts in the microbial community of control samples (IC and OC) from April–August and December. The relative abundance of *Chlaymdiae* (Family: *Simkaniaceae*), the second abundant phylum in the outer Bay (OC) with all zOTUs (10 in count) belonging to “Unclassified Simakaniaceae,” decreased from April to August before increasing slightly in December. *Tenericutes* (specifically *Mollicutes*), the second dominant phylum in the Inner Bay control samples (IC), increased in abundance over time, peaking in December. One zOTU annotated as “Unclassified Entomoplasmatales” had the highest abundance among different members of *Tenericutes*, including zOTUs belonging to *Acholeplasma*, *Mycoplasma*, and *Candidatus* (Bacilloplasma), and *Candidatus* (Hepatoplasma). In July, we observed a sudden spike in *Verrucomicrobia* abundance in two samples from IC, but soon after in August the community composition became similar to that in June. We also observed patterns of community dynamics in *Chlamydiae* and *Tenericutes* in cross-transplant samples (I→O and O→I) over the sampling period. *Chlamydiae* became the most dominant group in O→I (May) samples, but its abundance decreased sharply thereafter, whereas in I→O samples, *Chlaymdiae* and *Tenericutes* both remained stable, with *Chlamydiae* being dominant in May and June, and *Tenericutes* being dominant in July and August (Figure 2A, Figure S2).

At the genus taxonomic rank, *Endozoicomonas* species were the most dominant. Forty-eight zOTUs (out of 2,064) were taxonomically classified as *Endozoicomonas*. These 48 zOTUs accounted for an average of ∼54% relative abundance in corals fragments and 0% in sea-water samples. Of these 48 zOTUs, 13 contributed ∼90% of the total *Endozoicomonas* abundance. Interestingly, IC and OC samples harbored different dominant *Endozoicomonas* zOTUs, with zOTU1 and zOTU2 being dominant in OC and zOTU7 and zOTU9 in IC. Across the sampling time, we also observed shifts in the dominant *Endozoicomonas* phylotypes, zOTU2 was dominant from April to June, whereas zOTU1 became dominant in July–December OC samples. In IC samples, zOTU7 was dominant from April to May, but after that its relative abundance declined (Figure 2B). It is also worth noting that a significant decline in the *Endozoicomonas* abundance was observed in IC samples in July, August, and December (Figure 2B), suggestive of a locational dependence.

In cross-transplant samples, the *Endozoicomonas* phylotypes from OC remained resistant to change when transplanted in the Inner Bay (O→I), with zOTU2 being dominant across all sampling times. For I→O transplanted samples, however, instead of *Endozoicomonas* phylotypes from IC, we observed that zOTU1—another dominant *Endozoicomonas* phylotype in OC samples—was dominant (Figure 2B), suggesting that the phylotypes had different robustnesses under different environmental scale disturbances.

### Location-dependent robustness in the coral microbiome

The dispersion of homogeneity analysis identified that the bacterial community in corals with the Inner Bay as the final location (IC and O→I) were significantly different from each other (ANOVA, F=9.23, *p* <0.001), whereas samples whose final destination was the Outer Bay (OC and I→O) had no significant difference (ANOVA, F=1.98, *p* >0.05), indicating that the microbiome had location specificity. Therefore, samples whose final destinations were the Inner Bay and Outer Bay were analyzed independently to test for differences in community composition between the control and transplant groups. Ordination analysis using nMDS, followed by PERMANOVA identified the significant influence of coral sample, sampling month, and their combined effect (interaction term) Figure 3A). Ellipses with a 95% confidence interval suggested that samples for which the Outer Bay was their final location (OC and I→O) were more similar to each other compared to samples for which the Inner Bay was their final location (IC and O→I). These findings support locational variability and differential robustness in the coral microbiome (Figure 3A–B). Transplanted samples (O→I) clustered tightly compared to IC samples, indicating less variability after transplantation in the transplant samples. However, highly overlapping ellipses were observed for OC and I→O samples, suggesting a highly similar microbial community in the control (OC) and transplanted samples from the Inner Bay (I→O) (Figure 3A–B).

**Figure 3.**
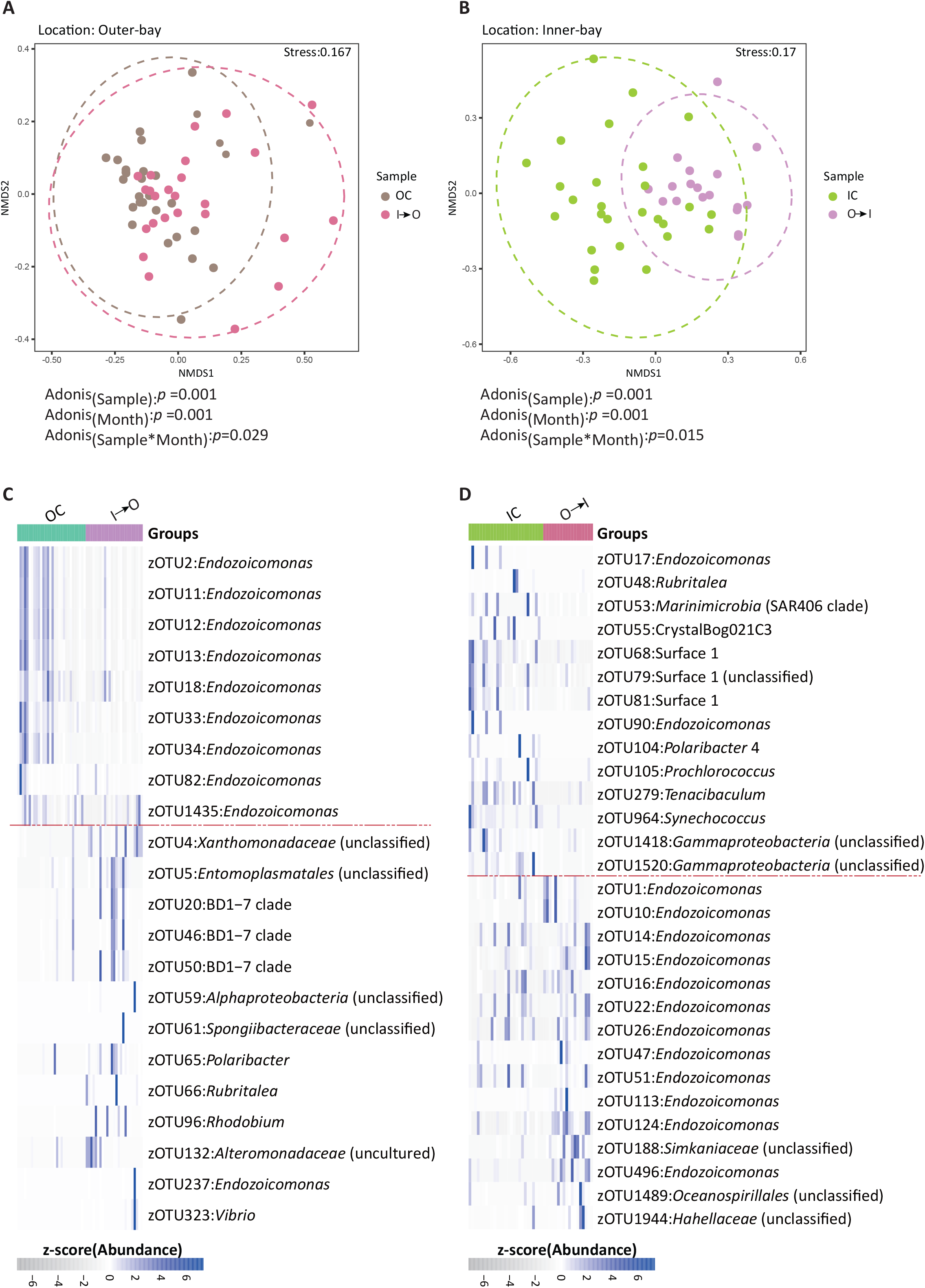
Location-dependent bacterial community structure and differentially abundant bacterial community. Plots based on non-metric multidimensional scaling (nMDS) of Bray-Curtis dissimilarity of bacterial community composition at the zOTUs level associated with different locations. Final location: **A)** Outer Bay (OC and I→O) and **B)** Inner Bay (IC and O→I). PERMANOVA analysis-identified sample location, month, and interaction terms are significant factors in determining the *Acropora muricata* microbiome. LefSe results based differentially abundant zOTUs over sample groups in the **C)** Outer Bay (OC and I→O) and **D)** Inner Bay (IC and O→I). zOTUs above the dotted red lines are differentially abundant in control (OC and IC) and the ones below are differentially abundant in transplant samples (I→O and O→I)

### Differentially abundant microbiome dominated by *Endozoicomonas*-related phylotypes

LEfSe analysis identified differentially abundant zOTUs of different taxa across all the sampling groups: nine zOTUs were differentially abundant in OC samples, 13 in I→O samples, and 16 in IC and O→I samples (Figure 3C–D). Interestingly, all the differentially abundant zOTUs in the OC samples belonged to *Endozoicomonas,* but zOTUs belonging to diverse taxa—including the BD1-7 clade (*Gammaproteobacteria*), *Entomoplasmatales* (Phylum: *Tenericutes*; Class: *Mollicutes*), and *Alteromonadaceae* (Class: *Gammaproteobacteria*)—were differentially abundant in I→O samples (Figure 3C). Similarly, out of the 16 zOTUs that were differentially abundant in O→I samples, 13 were *Endozoicomonas*; IC samples also had zOTUs belonging to diverse taxa that were differentially abundant, including Surface 1_ge (Class: *Alphaproteobacteria*), *Synechococcus* (Class: *Cyanobacteria*), and others (Figure 3D).

### Phylogenetic analysis of dominant *Endozoicomonas* zOTUs and a novel cultured species

The high abundance of *Endozoicomonas*-related phylotypes in the coral samples and their differential robustness after transplantation a) motivated us to determine their phylogenetic position and b) provided an opportunity to isolate and culture these phylotypes. A phylogenetic tree based on 16S rRNA gene sequences and the percentage identity (% identity) match between these sequences confirmed that zOTU1 and zOTU2 were 99.02 and 98.05% identical (16S rRNA V6-V8 region), respectively, to *Endozoicomonas acroporae* Acr-14^T^ (Figure 4A). They also formed a distinct clade with zOTU10, zOTU13, zOTU15, and zOTU18 (Figure 4A). These zOTUs (zOTU 10, 13,15, and 18) were also >97% identical to *E. acroporae* Acr-14^T^ 16S rRNA gene (Figure S3A). However, zOTU7, zOTU9, zOTU16, and zOTU17 formed a separate clade away from any cultured *Endozoicomonas* species (Figure 4A). zOTU7 was 100% identical to a newly isolated and cultured species (*Ca.* E. penghunesis 4G) described in this study (see sections below) and zOTU9 had 98.70% identity (16S rRNA gene V6-V8 region) with *Ca.* E. penghunesis 4G (Figure 4A). zOTU17 and zOTU16 were also >97% identical to *Ca.* E. penghunesis 4G 16S rRNA gene (Figure S3B). A genomic analysis of *Ca.* E. penghunesis 4G identified seven copies of 16S rRNA (see the section below) based on percentage similarity; 16S rRNA gene copy (1) (Figure S3C) was used as the representative for the phylogenetic tree in Figure 4A. We also performed phylogenetic analysis for all copies of 16S rRNA present in *Ca.* E. penghunesis and *E. acroporae* Acr-14^T^ (Figure S3 D).

**Figure 4.**
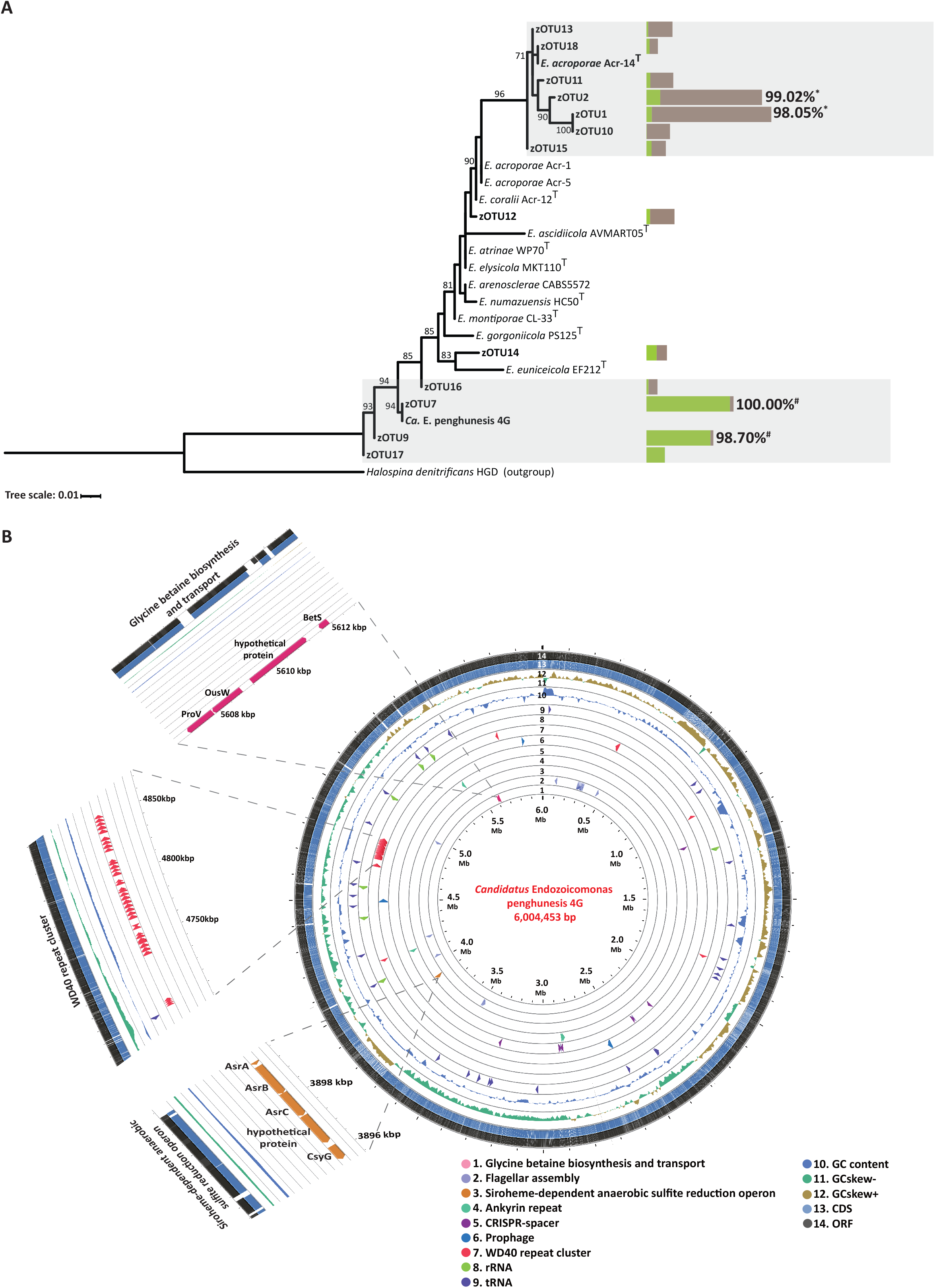
Phylogenetic tree and genome map of *Candidatus* Endozoicomonas penghunesis 4G. **A)** Phylogenetic tree of dominant zOTUs and 16S rRNA sequences of *Ca*. E. penghunesis (Copy1) and *Endozoicomonas acroporae* Acr-14^T^. Horizontal bars denote the relative abundance of selective zOTUs in the Inner (Green) and Outer Bay (Brown). The percent values denote the percentage identity between the zOTU and cultured 16S rRNA copy, with “#” corresponding to *Ca.* E. penghunesis and “*” to *E. acroporae* Acr-14^T^. Shaded regions are considered to belong to one bacterial species. **B)** Whole-genome map of *Ca.* E. penghunesis 4G drawn in CGViewer with concentric circles depicting distinct features. The map also highlights the concentration of WD40 domain proteins, Siroheme-dependent anaerobic sulfite reduction operon, and Glycine-betaine biosynthesis and transport pathways.

### Description of *Ca.* Endozoicomonas penghunesis 4G

*Ca.* E. penghunesis 4G is a gram-negative, facultatively anaerobic, and slightly motile bacterium that forms beige-colored colonies (size=2.14×0.66 µm) and is slightly susceptible to the antibiotics streptomycin and ampicillin. No catalase enzymatic activity was reported for this bacterium, but the bacterial culture was trypsin and oxidase-positive (Table S2). This new bacterial species tolerates the widest temperature (15–35°C) and salinity (5–30 PSU) ranges of the characterized *Endozoicomonas* species (Table S3). The pH range for growth was pH 6.0–10.0, with optimal growth observed at a slightly alkaline pH (pH 8.0). SEM and TEM analyses identified rod-shaped cells (Figure S4C) surrounded by a possible mucus lining (Figure S4A) and with structures that appeared to be granules or vacuoles in the cell (Figure S4B). A 16S rRNA gene sequence blast search identified the closest cultured relative to be *Endozoicomonas montiporae* CL-33 (Accession ID: CP013251) with 96.17% identity; based on a species identity cutoff of 97%, this suggests that the bacterium is a novel *Endozoicomonas* species.

### Genome assembly features of *Ca.* E. penghunesis 4G

The genome of *Ca.* E. penghunesis 4G was first assembled using Nanopore reads and later polished with quality filtered Illumina reads, resulting in a single contig of 6,004,453 bp and N50 (6,004,453). Genome completeness, contamination, and strain heterogeneity were estimated to be 97.52, 0.98, and 0%, respectively. Out of 573 single-copy marker genes (c_Gammaproteobacteria) from the checkM database [61], 493 genes were present only once, five single-copy markers were duplicated, and nine were missing. Based on the criteria of “minimum information for single amplified and metagenome-assembled genome of bacteria” [69], our genome can be considered “finished.” The GC content of the genome was 49.1%, which is similar to that of other *Endozoicomonas* species.

### Genomic features of *Ca.* E. penghunesis 4G

A total of 5,019 genes and 4,913 CDS were predicted from the genome. We annotated seven copies of 16S, nine of 23S, eight of 5S rRNA genes, and 80 tRNAs. A sequence similarity analysis of 16S rRNA gene copies revealed that all copies were at least 98.76% identical, with four copies >99.28% identical and two 100% identical to each other (Figure S3C). Copy-1 of the 16S rRNA gene was used as a representative sequence to classify the closest relative of dominant *Endozoicomonas* zOTUs identified in this study (Figure 4A). There were no CRISPR elements and only one prophage was identified in the genome. Out of the 5,019 genes predicted, more than 50% (2,721) were annotated to be hypothetical. Since, *Endozoicomonas* species have been exclusively isolated from their marine eukaryotic hosts, including *Ca.* E. penghunesis 4G, we searched for ELPs and identified 43 WD40 domain proteins (WD40), four Ankyrin repeat proteins (ARPs), and 12 Tetratricopeptide repeat proteins (TRPs). Almost all the WD40 domain-containing proteins were arranged consecutively (Figure 4B) and flanked by transposes. Most (27 out of 41) of the WD40 domain proteins were annotated as TolB protein from the Tol-Pal system, which is important for maintaining cellular integrity, and others (14) were classified as hypothetical proteins. A wide array of secretory proteins were also annotated with 248 Type III secretion system effectors (T3SS), 50 Type IV secretion system effectors (T4SS), and 10 Type VI secretion system effectors (T6SS) annotated from the proteome.

### Metabolic repertoire of *Ca.* E. penghunesis 4G

RAST classified only 36% (1,756) of the total genes into subsystems. Subsystems a) Carbohydrates and b) Cofactors, Vitamins, Prosthetic groups, and Pigments had the highest number of annotated genes—270 and 238, respectively (Figure S5). In the stress response subsystem, 108 genes were annotated, most of which were related to oxidative stress response (46 genes), followed by heat shock response (18) and detoxification response (16). Interestingly, within the osmotic stress response, we identified genes for betaine transport via ATP-binding cassette transporter, BetS (high-affinity choline uptake protein BetS), arranged in tandem with an L-proline glycine betaine ABC transport system permease (ProV and OusW) (Figure 4B). Multiple copies of superoxide dismutase, alkyl hydroperoxide reductase, and methionine sulfoxide reductase genes were also identified in the genome.

*Ca.* E. penghunesis 4G had genes encoding essential amino acids and pathways including glycolysis and tricarboxylic acid cycle; genes for converting nitrate to nitrite (NapAB, KO: K02567) and ammonia to L-glutamate were identified, but none related to the conversion of nitrite to ammonia. Assimilatory and dissimilatory sulfate reduction and oxidation pathways were also completely absent. Furthermore, no genes related to the uptake of extracellular taurine or its metabolism to sulfite were identified. Interestingly, siroheme biosynthesis and siroheme-dependent anaerobic sulfite reduction operons were present in *Ca.* E. penghunesis 4G (Figure 4B). We also identified genes arranged in an operon-like manner for anaerobic glycerol degradation.

### Comparing *E. acroporae* and *Ca.* E. penghunesis physiological and genomic features

*Endozoicomonas* phylotypes belonging to *E. acroporae* and *Ca*. E. penghunesis 4G were dominant in colonies of coral *Acropora muricata* in the Outer and Inner Bay, respectively (figure 2B, 4A). This selective dominance could be attributed to multiple factors, including bacterial physiological and genetic repertoire. Therefore, we compared the two species—*Ca.* E. penghunesis had a wider growth temperature range compared to *E. acroporae* and was slightly motile. *E. acroporae*, on the other hand, was non-motile (Table S2) and had a wider salinity and growth pH range (Table S3). A wider growth temperature range of *Ca.* E. penghunesis and its dominance in the Inner Bay aligns with the high variation of temperature fluctuations (from summer to winter) observed in the calmer waters of the Inner Bay. Comparing the metabolic repertoire of the two species, we found that genes for dimethylsulfoniopropionate (DMSP) metabolism and dimethylsulfoxide (DMSO) reduction were absent from *Ca.* E. penghunesis, but *E. acroporae* had a complete operon for DMSP metabolism as reported in our previous study [33] as well as genes for DMSO reduction. Lack of a potent oxidative stress response gene repertoire potential could be the reason for the loss of abundance of *Ca.* E. penghunesis in the control (IC) and transplant samples (I→O) in summer (June-August). Furthermore, the robustness of *E. acroporae* in the control (OC) and transplant samples (O→I) throughout the year could be due to the species’ ability to remove oxidative stress (which increased in summer) more efficiently via DMSP metabolism and the presence of catalase activity.

### Coral mortality in the Inner Bay

We observed coral mortality exclusively in the Inner Bay for both IC and O→I samples (marked as ‘X’, Figure 2A-B). Grazing by *Drupella cornus* was also observed in the samples from the Inner Bay during the sampling in May, and continued until December (Figure S6).

## Discussion

This study aimed to test the differential adaptation capability of *Endozoicomonas,* one of the most dominant bacterial groups in the coral microbiome. We analyzed the microbiome dynamics of the common Indo-Pacific coral *A. muricata* over 9 months following a reciprocal transplant experiment at the finest resolution of zOTUs. We identified that different *Endozoicomonas* phylotypes in the coral *A. muricata* coral colonies belonging to two dominant species, including a novel species, have differential adaptation capabilities, with one species more resilient to change than the other. Our results shed light on an often-neglected factor when determining variations in community composition: bacterial species/strains adapt differently when coral hosts are subjected to biotic and abiotic stressors. Furthermore, we also isolated, cultured, and sequenced a single chromosome-level genome of one of the dominant phylotypes belonging to a novel *Endozoicomonas* species, *Ca.* Endozoicomonas penghunesis 4G, to ascertain the ecological and functional role of this bacterium and add to the growing knowledge and genome datasets of this key microbe in coral reefs.

*Acropora muricata* microbiome was dominated by members of class *Gammaproteobacteria* (Phylum: *Proteobacteria*) (Silva v132 grouped *Gammaproteobacteria* into *Betaproteobacteria*), particularly by *Endozoicomonas-*related phylotype*s* (Figure 2A–B; Figure S2). Members of genus *Endozoicomonas* are often found to be the dominant group in the microbiome of several coral species—e.g., members of *Acropora*, *Pocillopora*, and *Stylophora* [3, 24, 30, 70]—and have been proposed to play a significant role in coral health and protection [2, 71] and coral sulfur cycling [33, 72]. Another dominant bacterial group*, Simkaniaceae* (Phylum: *Chlamydiae*, Class: *Chlamydiae*), was described as an obligate intracellular bacterium, but its function has remained enigmatic [70, 73]. Like *Endozoicomonas*, *Simkaniaceae*-related phylotypes were also recently found to be abundant in healthy corals from the reefs in Florida, but their abundance decreased in corals suffering from stony coral tissue loss disease [74]. Members of class *Mollicutes—*particularly zOTUs related to *Entomoplasmatales* and *Mycoplasmatales—*are suggested to be mutualistic or commensal bacteria in temperate and deep-sea gorgonians and cold-water Scleractinia corals, but their specific function remains unknown [75, 76]. Overall, the microbial community composition in the coral colonies of the control group remained stable throughout the experiment timeline, with only transient differences observed between them across sampling time (Fig 2A and 2B).

Spatial and temporal fluctuations in the microbiome were observed throughout the experiment, community abundance (Figure 2A) and ordination analysis showed that community structure varies across temporal and spatial scales (Figure 3A). Varying degrees of overlap between the samples from the same location suggest that the microbes show different scales of variability, and that microbial structure is a function of the local environment. Previous studies have also shown that the microbiome varies spatially due to differences in the sites’ local environments [18, 77]. Ocean currents are believed to have a homogenizing effect on the microbial communities, and the same coral species separated by hundreds to thousands of kilometers have been found to have similar microbiome compositions [1, 4]. Our results were surprising in this regard, as a high site-to-site variation was observed at a relatively small scale. A potential reason for this variation could be high abiotic and anthropogenic pollution in the Inner Bay compared to the Outer bay; similar results were obtained in an earlier study by Ziegler et al [27].

Utilizing the zOTU approach for metabarcoding data analysis and focusing on the dominant bacterial genus, we identified that colonies of *A. muricata* from the Inner and Outer Bay were not dominated by a single *Endozoicomonas* phylotype, but had several differentially abundant phylotypes associated with them (Figure 2B). These results are similar to the multiple phylotypes that were reported to be dominant in colonies of the same coral species as identified earlier [70]. However, the dominant *Endozoicomonas* phylotypes identified in our study were different in coral colonies from the two locations, with Inner Bay colonies harboring a novel species *Ca.* E. penghunesis and Outer Bay colonies harboring *E. acroporae-*related phylotypes (Figure 4A). This result was intriguing, as corals of the *Acropora* genus are known to have a strong influence on their microbial community composition [18]. Another observation that arises from these results is how the single-nucleotide variation approach utilized to obtain ASVs or zOTUs can potentially lead to increased diversity (richness) estimates, which rely on ASV or zOTU counts, especially with bacterial groups known to have more than one copy of non-identical 16S rRNA genes in their genome. In our study this is true in the case of the genus *Endozoicomonas*: members of this genus are known to harbor more than one copy of 16S rRNA, complete genomes of *Endozoicomonas montiporae* CL-33^T^ have seven copies [52], similar to *Ca.* E. penghunesis, that are not all identical (Figure S3); hence, we used a phylogenetic approach to assign the taxonomy to these *Endozoicomonas* phylotypes.

Members of the coral holobiont potentially engage in complex interactions to maintain the health and fitness of the coral host, and external stressors could disturb these interactions by influencing the composition of the holobiont. To overcome the influence of the external stressors, a genomic adaptation of the holobiont members (in our case, bacteria) could play an important role in host survival. Since, a bacterium is likely to be not as well adapted to a new niche as resident strains, unless it has the genetic capability to mitigate the new stressors [78, 79]. During our reciprocal transplant experiment, we observed that *E. acroporae*-related zOTUs remained dominant in the control (OC) and the transplanted samples (O→I), with only change in the dominant zOTU from zOTU1 to zOTU2, whereas, those related to *Ca.* E. penghunesis 4G were only dominant in April and May months in the control samples of the Inner Bay (IC) (Figure 2B).

There are a few possible reasons for this observation. One is that *E. acroporae* may be more resilient and better adapted to diverse conditions encountered in the Inner and the Outer Bay compared to *Ca.* E. penghunesis 4G. However, further investigation is required to say this with greater confidence. In addition, variation in the abundance of the different *Endozoicomonas* phylotypes is potentially analogous to the abundance of genotypes of different endosymbiotic algae *Symbiodinium*, often found in coral colonies [80]. Several coral species can perform “symbiont shuffling” to select for the more thermotolerant genotype of endosymbiotic algae in response to thermal stress [81–83]. We observed microbial shuffling in *Endozoicomonas* phylotypes to a certain degree in our transplant samples, where zOTU2 became dominant in O→I samples and zOTU1 became dominant in I→O sample, although the genus’ abundance was very low in IC samples, which were dominated by zOTU7 and zOTU9 (Figure 2B). However, the selective advantages or potential benefits of shuffling microbiome members are unclear and require further explorations.

Underpinning the functional and ecological role of coral-associated microbes in reefs has become critical to developing an intervention for coral reef protection, such as developing a coral probiotic [2, 84]. These interventions require in-depth information about members of the coral holobiont. In the current study, we isolated, cultured, and sequenced the complete genome of a dominant *Endozoicomonas* phylotype identified in the metabarcoding data analysis. Phylogenetic analysis identified that the dominant zOTUs (zOTU1, zOTU2, zOTU10, zOTU11, zOTU13, and zOTU18) from the Outer Bay were closer to a previously characterized species *E. acroporae* [85], whose genome was sequenced earlier [33, 62].

On the other hand, the Inner Bay-dominant zOTUs (zOTU7, zOTU9, zOTU16, and zOTU17) were closest to the novel species *Ca.* E. penghunesis 4G isolated and characterized in this study. Genomic analysis of *Ca.* E. penghunesis revealed features similar to other *Endozoicomonas* species—i.e., large genome size (∼6.00 Mb), many coding genes (4,913), and complete pathways for essential amino acids—suggesting a free-living life stage, yet *Endozoicomonas* have been called an “obligate” endosymbiont of corals [71]. This assertion of an “obligate” endosymbiont stems from the almost negligible abundance of *Endozoicomonas* in the coral-ecosphere [86] and very low abundance in early life stages of coral [80, 87]. Furthermore, a comparative genomics study identified that *Endozoicomonas* species are capable of differential functional specificity, and different genotypes may play disparate metabolic roles in their hosts [34]. This is true for sulfur metabolism, where *E. acroporae* is the only known *Endozoicomonas* species capable of metabolizing dimethylsulfoniopropionate (DMSP) to dimethylsulfide (DMS) [33], genes for DMSP metabolism operon were not present in the genome of *Ca.* E. penghunesis. However, transporters (three copies) for glycine-betaine, another osmolyte, were identified in the genome of *Ca.* E. penghunesis. Other *Endozoicomonas* species have also been identified to have the ability to scavenge glycine-betaine through transporters [88], potentially to alleviate oxidative stress. Identification of putative siroheme-dependent anaerobic sulfite reduction operon was interesting as this process facilitates growth under anaerobic conditions (B_12_-dependent anaerobic growth) by oxidizing 1,2-propanediol with tetrathionate as an electron acceptor [89]. Physiological tests also showed that *Ca.* E. penghunesis is a facultative anaerobe; however, more functional evidence is required to confirm this outcome and the advantage (if any) it provides the bacterium and coral host that maintains it.

We observed coral mortality during our experiment exclusively in the Inner Bay, the corallivorous snail *Drupella cornus,* which exclusively feeds on living tissue, grazed there (Figure S6). These gastropods occur throughout the shallow waters of the Indo-Pacific region [90]. Outbreaks of this corallivorous marine gastropod have been recorded in different parts of the Gulf of Eilat, Israel [91] and the Great Barrier Reef, Australia [92]. Coral feeding gastropods of *Drupella* sp. show a strong preference for preying on *Acroporids* [93] and are known to be efficient vectors for brown band disease in corals [92, 94]. Though no *Drupella* sp. outbreaks to date have been recorded in Taiwan’s coral reefs and no visible signs of brown band disease were observed in our study, it is important to keep monitoring the corals in the Penghu Archipelago for signs of climate change and disease outbreaks in the near future.

## Conclusion

A variety of factors, many of which are external, are known to influence the coral microbiome composition and its dynamics. However, an important internal factor, the adaptation capability of microbiome members, which governs the survival of a bacterium in a niche, has been overlooked. Using a combination of metabarcoding, genomic, and comparative genomic approaches, we showed that members of the dominant bacterial group *Endozoicomonas* are capable of sustaining and proliferating in a new niche following a reciprocal transplant experiment. Our ability to isolate and culture one of the dominant bacterial species, *Ca.* Endozoicomonas penghunesis 4G, builds on our knowledge of these important bacterial groups in the coral holobiont. Furthermore, we address critical aspects of using zOTUs/ASVs to estimate bacterial richness using metabarcoding data, which can result in often falsely inflated diversity estimates, especially in the case of microbes harboring more than one copy of non-identical 16S rRNA gene, e.g. *Endozoicomonas*. In summary, we conclude that different members of the coral holobiont belonging to the same bacterial group can have differential adaptation capabilities, and this internal factor should also be considered when devising interventions to protect coral reefs, like developing a coral probiotic.

## Acknowledgement

This study was supported by funding to SLT from Academia Sinica and the Ministry of Science and Technology. We would like to thank Noah Last of Third Draft Editing for his English language editing.

## References

1. Rohwer F, Seguritan V, Azam F, Knowlton N. Diversity and distribution of coral-associated bacteria. Marine Ecology Progress Series . 2002., 243: 1–10

2. Peixoto RS, Rosado PM, Leite DC de A, Rosado AS, Bourne DG. Beneficial Microorganisms for Corals (BMC): Proposed Mechanisms for Coral Health and Resilience. Front Microbiol 2017; 8: 341.

3. van Oppen MJH, Blackall LL. Coral microbiome dynamics, functions and design in a changing world. Nat Rev Microbiol 2019; 17: 557–567.

4. Dinsdale EA, Pantos O, Smriga S, Edwards RA, Angly F, Wegley L, et al. Microbial ecology of four coral atolls in the Northern Line Islands. PLoS One 2008; 3: e1584.

5. Blackall LL, Wilson B, van Oppen MJH. Coral-the world’s most diverse symbiotic ecosystem. Mol Ecol 2015; 24: 5330–5347.

6. Bourne DG, Munn CB. Diversity of bacteria associated with the coral Pocillopora damicornis from the Great Barrier Reef. Environ Microbiol 2005; 7: 1162–1174.

7. Li J, Chen Q, Long L-J, Dong J-D, Yang J, Zhang S. Bacterial dynamics within the mucus, tissue and skeleton of the coral Porites lutea during different seasons. Sci Rep 2014; 4: 1–8.

8. Glasl B, Herndl GJ, Frade PR. The microbiome of coral surface mucus has a key role in mediating holobiont health and survival upon disturbance. ISME J 2016; 10: 2280–2292.

9. Yang S-H, Tseng C-H, Huang C-R, Chen C-P, Tandon K, Lee STM, et al. Long-Term Survey Is Necessary to Reveal Various Shifts of Microbial Composition in Corals. Front Microbiol 2017; 8: 1094.

10. Pollock FJ, McMinds R, Smith S, Bourne DG, Willis BL, Medina M, et al. Coral-associated bacteria demonstrate phylosymbiosis and cophylogeny. Nat Commun 2018; 9: 4921.

11. Agostini S, Suzuki Y, Higuchi T, Casareto BE, Yoshinaga K, Nakano Y, et al. Biological and chemical characteristics of the coral gastric cavity. Coral Reefs 2012; 31: 147–156.

12. Marcelino VR, van Oppen MJ, Verbruggen H. Highly structured prokaryote communities exist within the skeleton of coral colonies. ISME J 2018; 12: 300–303.

13. Yang S-H, Tandon K, Lu C-Y, Wada N, Shih C-J, Hsiao SS-Y, et al. Metagenomic, phylogenetic, and functional characterization of predominant endolithic green sulfur bacteria in the coral Isopora palifera. Microbiome 2019; 7: 3.

14. Ricci F, Fordyce A, Leggat W, Blackall LL, Ainsworth T, Verbruggen H. Multiple techniques point to oxygenic phototrophs dominating the Isopora palifera skeletal microbiome. Coral Reefs 2021; 40: 275–282.

15. Littman RA, Willis BL, Pfeffer C, Bourne DG. Diversities of coral-associated bacteria differ with location, but not species, for three acroporid corals on the Great Barrier Reef. FEMS Microbiol Ecol 2009; 68: 152–163.

16. Kvennefors ECE, Sampayo E, Ridgway T, Barnes AC, Hoegh-Guldberg O. Bacterial communities of two ubiquitous Great Barrier Reef corals reveals both site- and species-specificity of common bacterial associates. PLoS One 2010; 5: e10401.

17. Morrow KM, Moss AG, Chadwick NE, Liles MR. Bacterial associates of two Caribbean coral species reveal species-specific distribution and geographic variability. Appl Environ Microbiol 2012; 78: 6438–6449.

18. Dunphy CM, Gouhier TC, Chu ND, Vollmer SV. Structure and stability of the coral microbiome in space and time. Sci Rep 2019; 9: 6785.

19. Epstein HE, Smith HA, Cantin NE, Mocellin VJL, Torda G, van Oppen MJH. Temporal Variation in the Microbiome of Acropora Coral Species Does Not Reflect Seasonality. Front Microbiol 2019; 10: 1775.

20. Bourne D, Iida Y, Uthicke S, Smith-Keune C. Changes in coral-associated microbial communities during a bleaching event. ISME J 2008; 2: 350–363.

21. Zaneveld JR, Burkepile DE, Shantz AA, Pritchard CE, McMinds R, Payet JP, et al. Overfishing and nutrient pollution interact with temperature to disrupt coral reefs down to microbial scales. Nat Commun 2016; 7: 11833.

22. Ziegler M, Seneca FO, Yum LK, Palumbi SR, Voolstra CR. Bacterial community dynamics are linked to patterns of coral heat tolerance. Nat Commun 2017; 8: 14213.

23. Shiu J-H, Keshavmurthy S, Chiang P-W, Chen H-J, Lou S-P, Tseng C-H, et al. Dynamics of coral-associated bacterial communities acclimated to temperature stress based on recent thermal history. Sci Rep 2017; 7: 14933.

24. Maher RL, Schmeltzer ER, Meiling S, McMinds R, Ezzat L, Shantz AA, et al. Coral Microbiomes Demonstrate Flexibility and Resilience Through a Reduction in Community Diversity Following a Thermal Stress Event. Frontiers in Ecology and Evolution 2020; 8: 356.

25. Wang L, Shantz AA, Payet JP, Sharpton TJ, Foster A, Burkepile DE, et al. Corals and their microbiomes are differentially affected by exposure to elevated nutrients and a natural thermal anomaly. Front Mar Sci 2018; 5.

26. Gignoux-Wolfsohn SA, Aronson FM, Vollmer SV. Complex interactions between potentially pathogenic, opportunistic, and resident bacteria emerge during infection on a reef-building coral. FEMS Microbiol Ecol 2017; 93.

27. Ziegler M, Roik A, Porter A, Zubier K, Mudarris MS, Ormond R, et al. Coral microbial community dynamics in response to anthropogenic impacts near a major city in the central Red Sea. Mar Pollut Bull 2016; 105: 629–640.

28. Osman EO, Suggett DJ, Voolstra CR, Pettay DT, Clark DR, Pogoreutz C, et al. Coral microbiome composition along the northern Red Sea suggests high plasticity of bacterial and specificity of endosymbiotic dinoflagellate communities. Microbiome 2020; 8: 8.

29. La Rivière M, Garrabou J, Bally M. Evidence for host specificity among dominant bacterial symbionts in temperate gorgonian corals. Coral Reefs 2015; 34: 1087–1098.

30. Ziegler M, Grupstra CGB, Barreto MM, Eaton M, BaOmar J, Zubier K, et al. Coral bacterial community structure responds to environmental change in a host-specific manner. Nat Commun 2019; 10: 3092.

31. Glasl B, Smith CE, Bourne DG, Webster NS. Disentangling the effect of host-genotype and environment on the microbiome of the coral Acropora tenuis. PeerJ 2019; 7: e6377.

32. Neave MJ, Rachmawati R, Xun L, Michell CT, Bourne DG, Apprill A, et al. Differential specificity between closely related corals and abundant Endozoicomonas endosymbionts across global scales. ISME J 2017; 11: 186–200.

33. Tandon K, Lu C-Y, Chiang P-W, Wada N, Yang S-H, Chan Y-F, et al. Comparative genomics: Dominant coral-bacterium Endozoicomonas acroporae metabolizes dimethylsulfoniopropionate (DMSP). ISME J 2020; 14: 1290–1303.

34. Neave MJ, Michell CT, Apprill A, Voolstra CR. Endozoicomonas genomes reveal functional adaptation and plasticity in bacterial strains symbiotically associated with diverse marine hosts. Sci Rep 2017; 7: 1–12.

35. Ribas-Deulofeu L, Denis V, De Palmas S, Kuo C-Y, Hsieh HJ, Chen CA. Structure of Benthic Communities along the Taiwan Latitudinal Gradient. PLoS One 2016; 11: e0160601.

36. Hsieh HJ, Hsien Y-L, Jeng M-S, Tsai W-S, Su W-C, Chen CA. Tropical fishes killed by the cold. Coral Reefs 2008; 27: 599–599.

37. Wilson K. Preparation of genomic DNA from bacteria. Curr Protoc Mol Biol 2001; Chapter 2: Unit 2.4.

38. Chen C-P, Tseng C-H, Chen CA, Tang S-L. The dynamics of microbial partnerships in the coral Isopora palifera. ISME J 2011; 5: 728–740.

39. Jorgensen SL, Hannisdal B, Lanzén A, Baumberger T, Flesland K, Fonseca R, et al. Correlating microbial community profiles with geochemical data in highly stratified sediments from the Arctic Mid-Ocean Ridge. Proc Natl Acad Sci U S A 2012; 109: E2846–55.

40. Edgar RC. UPARSE: highly accurate OTU sequences from microbial amplicon reads. Nat Methods 2013; 10: 996–998.

41. Schloss PD, Westcott SL, Ryabin T, Hall JR, Hartmann M, Hollister EB, et al. Introducing mothur: open-source, platform-independent, community-supported software for describing and comparing microbial communities. Appl Environ Microbiol 2009; 75: 7537–7541.

42. Edgar RC, Haas BJ, Clemente JC, Quince C, Knight R. UCHIME improves sensitivity and speed of chimera detection. Bioinformatics 2011; 27: 2194–2200.

43. Edgar RC. UNOISE2: improved error-correction for Illumina 16S and ITS amplicon sequencing. bioRxiv. 2016., 081257

44. Quast C, Pruesse E, Yilmaz P, Gerken J, Schweer T, Yarza P, et al. The SILVA ribosomal RNA gene database project: improved data processing and web-based tools. Nucleic Acids Res 2013; 41: D590–6.

45. Yilmaz P, Parfrey LW, Yarza P, Gerken J, Pruesse E, Quast C, et al. The SILVA and “All-species Living Tree Project (LTP)” taxonomic frameworks. Nucleic Acids Res 2014; 42: D643–8.

46. McMurdie PJ, Holmes S. phyloseq: an R package for reproducible interactive analysis and graphics of microbiome census data. PLoS One 2013; 8: e61217.

47. Oksanen J, Kindt R, Legendre P, O’Hara B, Simpson GL, Solymos P, et al. vegan: Community Ecology Package. 2008.

48. Wickham H. ggplot2. Wiley Interdiscip Rev Comput Stat 2011; 3: 180–185.

49. Kolde R. pheatmap: Pretty heatmaps. Github.

50. Cao Y. microbiomeMarker: R package for microbiome biomarker discovery. Github.

51. Shiu J-H, Ding J-Y, Tseng C-H, Lou S-P, Mezaki T, Wu Y-T, et al. A Newly Designed Primer Revealed High Phylogenetic Diversity of Endozoicomonas in Coral Reefs. Microbes Environ 2018; 33: 172–185.

52. Ding J-Y, Shiu J-H, Chen W-M, Chiang Y-R, Tang S-L. Genomic Insight into the Host-Endosymbiont Relationship of Endozoicomonas montiporae CL-33(T) with its Coral Host. Front Microbiol 2016; 7: 251.

53. Nawrocki EP, Eddy SR. Infernal 1.1: 100-fold faster RNA homology searches. Bioinformatics 2013; 29: 2933–2935.

54. Kalvari I, Argasinska J, Quinones-Olvera N, Nawrocki EP, Rivas E, Eddy SR, et al. Rfam 13.0: shifting to a genome-centric resource for non-coding RNA families. Nucleic Acids Res 2018; 46: D335–D342.

55. Trifinopoulos J, Nguyen L-T, von Haeseler A, Minh BQ. W-IQ-TREE: a fast online phylogenetic tool for maximum likelihood analysis. Nucleic Acids Res 2016; 44: W232–5.

56. Letunic I, Bork P. Interactive Tree Of Life (iTOL) v4: recent updates and new developments. Nucleic Acids Res 2019; 47: W256–W259.

57. De Coster W, D’Hert S, Schultz DT, Cruts M, Van Broeckhoven C. NanoPack: visualizing and processing long-read sequencing data. Bioinformatics 2018; 34: 2666–2669.

58. Kolmogorov M, Bickhart DM, Behsaz B, Gurevich A, Rayko M, Shin SB, et al. metaFlye: scalable long-read metagenome assembly using repeat graphs. Nat Methods 2020; 17: 1103–1110.

59. Schubert M, Lindgreen S, Orlando L. AdapterRemoval v2: rapid adapter trimming, identification, and read merging. BMC Res Notes 2016; 9: 88.

60. Walker BJ, Abeel T, Shea T, Priest M, Abouelliel A, Sakthikumar S, et al. Pilon: an integrated tool for comprehensive microbial variant detection and genome assembly improvement. PLoS One 2014; 9: e112963.

61. Parks DH, Imelfort M, Skennerton CT, Hugenholtz P, Tyson GW. CheckM: assessing the quality of microbial genomes recovered from isolates, single cells, and metagenomes. PeerJ . 2015.

62. Tandon K, Chiang P-W, Chen W-M, Tang S-L. Draft Genome Sequence of Endozoicomonas acroporae Strain Acr-14T, Isolated from Acropora Coral. Genome Announc 2018; 6.

63. Seemann T. Prokka: rapid prokaryotic genome annotation. Bioinformatics 2014; 30: 2068–2069.

64. Aziz RK, Bartels D, Best AA, DeJongh M, Disz T, Edwards RA, et al. The RAST Server: rapid annotations using subsystems technology. BMC Genomics 2008; 9: 75.

65. Kanehisa M, Sato Y, Morishima K. BlastKOALA and GhostKOALA: KEGG tools for functional characterization of genome and metagenome sequences. J Mol Biol 2016; 428: 726–731.

66. Couvin D, Bernheim A, Toffano-Nioche C, Touchon M, Michalik J, Néron B, et al. CRISPRCasFinder, an update of CRISRFinder, includes a portable version, enhanced performance and integrates search for Cas proteins. Nucleic Acids Res 2018; 46: W246–W251.

67. Yang M, Derbyshire MK, Yamashita RA, Marchler-Bauer A. NCBI’s Conserved Domain Database and tools for protein domain analysis. Curr Protoc Bioinformatics 2020; 69: e90.

68. Grant JR, Stothard P. The CGView Server: a comparative genomics tool for circular genomes. Nucleic Acids Res 2008; 36: W181–4.

69. Bowers RM, The Genome Standards Consortium, Kyrpides NC, Stepanauskas R, Harmon-Smith M, Doud D, et al. Minimum information about a single amplified genome (MISAG) and a metagenome-assembled genome (MIMAG) of bacteria and archaea. Nat Biotechnol 2017; 35: 725–731.

70. Damjanovic K, Blackall LL, Peplow LM, van Oppen MJH. Assessment of bacterial community composition within and among Acropora loripes colonies in the wild and in captivity. Coral Reefs 2020; 39: 1245–1255.

71. Neave MJ, Apprill A, Ferrier-Pagès C, Voolstra CR. Diversity and function of prevalent symbiotic marine bacteria in the genus Endozoicomonas. Appl Microbiol Biotechnol 2016; 100: 8315–8324.

72. Raina J-B, Tapiolas D, Willis BL, Bourne DG. Coral-associated bacteria and their role in the biogeochemical cycling of sulfur. Appl Environ Microbiol 2009; 75: 3492–3501.

73. Collingro A, Tischler P, Weinmaier T, Penz T, Heinz E, Brunham RC, et al. Unity in variety--the pan-genome of the Chlamydiae. Mol Biol Evol 2011; 28: 3253–3270.

74. Meyer JL, Castellanos-Gell J, Aeby GS, Häse CC, Ushijima B, Paul VJ. Microbial Community Shifts Associated With the Ongoing Stony Coral Tissue Loss Disease Outbreak on the Florida Reef Tract. Front Microbiol 2019; 10: 2244.

75. Gray MA, Stone RP, McLaughlin MR, Kellogg CA. Microbial consortia of gorgonian corals from the Aleutian islands. FEMS Microbiol Ecol 2011; 76: 109–120.

76. van de Water JAJM, Melkonian R, Voolstra CR, Junca H, Beraud E, Allemand D, et al. Comparative Assessment of Mediterranean Gorgonian-Associated Microbial Communities Reveals Conserved Core and Locally Variant Bacteria. Microb Ecol 2017; 73: 466–478.

77. Hernandez-Agreda A, Leggat W, Bongaerts P, Ainsworth TD. The Microbial Signature Provides Insight into the Mechanistic Basis of Coral Success across Reef Habitats. MBio 2016; 7.

78. Hibbing ME, Fuqua C, Parsek MR, Peterson SB. Bacterial competition: surviving and thriving in the microbial jungle. Nat Rev Microbiol 2010; 8: 15–25.

79. Sheppard SK, Guttman DS, Fitzgerald JR. Population genomics of bacterial host adaptation. Nat Rev Genet 2018; 19: 549–565.

80. Quigley KM, Davies SW, Kenkel CD, Willis BL, Matz MV, Bay LK. Deep-sequencing method for quantifying background abundances of symbiodinium types: exploring the rare symbiodinium biosphere in reef-building corals. PLoS One 2014; 9: e94297.

81. Cunning R, Silverstein RN, Baker AC. Investigating the causes and consequences of symbiont shuffling in a multi-partner reef coral symbiosis under environmental change. Proceedings of the Royal Society B: Biological Sciences 2015; 282: 20141725.

82. Bay LK, Doyle J, Logan M, Berkelmans R. Recovery from bleaching is mediated by threshold densities of background thermo-tolerant symbiont types in a reef-building coral. R Soc Open Sci 2016; 3: 160322.

83. Boulotte NM, Dalton SJ, Carroll AG, Harrison PL, Putnam HM, Peplow LM, et al. Exploring the Symbiodinium rare biosphere provides evidence for symbiont switching in reef-building corals. ISME J 2016; 10: 2693–2701.

84. Peixoto RS, Sweet M, Villela HDM, Cardoso P, Thomas T, Voolstra CR, et al. Coral Probiotics: Premise, Promise, Prospects. Annu Rev Anim Biosci 2021; 9: 265–288.

85. Sheu S-Y, Lin K-R, Hsu M-Y, Sheu D-S, Tang S-L, Chen W-M. Endozoicomonas acroporae sp. nov., isolated from Acropora coral. Int J Syst Evol Microbiol 2017; 67: 3791–3797.

86. Weber L, Gonzalez-Díaz P, Armenteros M, Apprill A. The coral ecosphere: A unique coral reef habitat that fosters coral–microbial interactions. Limnol Oceanogr 2019; 64: 2373–2388.

87. Lema KA, Bourne DG, Willis BL. Onset and establishment of diazotrophs and other bacterial associates in the early life history stages of the coral Acropora millepora. Mol Ecol 2014; 23: 4682–4695.

88. Ngugi DK, Ziegler M, Duarte CM, Voolstra CR. Genomic Blueprint of Glycine Betaine Metabolism in Coral Metaorganisms and Their Contribution to Reef Nitrogen Budgets. iScience 2020; 23: 101120.

89. Price-Carter M, Tingey J, Bobik TA, Roth JR. The alternative electron acceptor tetrathionate supports B12-dependent anaerobic growth of Salmonella enterica serovar typhimurium on ethanolamine or 1,2-propanediol. J Bacteriol 2001; 183: 2463–2475.

90. Claremont M, Reid DG, Williams ST. Evolution of corallivory in the gastropod genus Drupella. Coral Reefs 2011; 30: 977–990.

91. Shafir S, Gur O, Rinkevich B. A Drupella cornus outbreak in the northern Gulf of Eilat and changes in coral prey. Coral Reefs 2008; 27: 379–379.

92. Nicolet KJ, Hoogenboom MO, Gardiner NM, Pratchett MS, Willis BL. The corallivorous invertebrate Drupella aids in transmission of brown band disease on the Great Barrier Reef. Coral Reefs 2013; 32: 585–595.

93. Moerland MS, Scott CM, Hoeksema BW. Prey selection of corallivorous muricids at Koh Tao (Gulf of Thailand) four years after a major coral bleaching event. Contrib Zool 2016; 85: 291–309.

94. Nicolet KJ, Chong-Seng KM, Pratchett MS, Willis BL, Hoogenboom MO. Predation scars may influence host susceptibility to pathogens: evaluating the role of corallivores as vectors of coral disease. Sci Rep 2018; 8: 5258.

