## Supplementary figures and images for "Microbiome restructuring: dominant coral bacterium *Endozoicomonas* species display differential adaptive capabilities to environmental changes"

### Figure S1

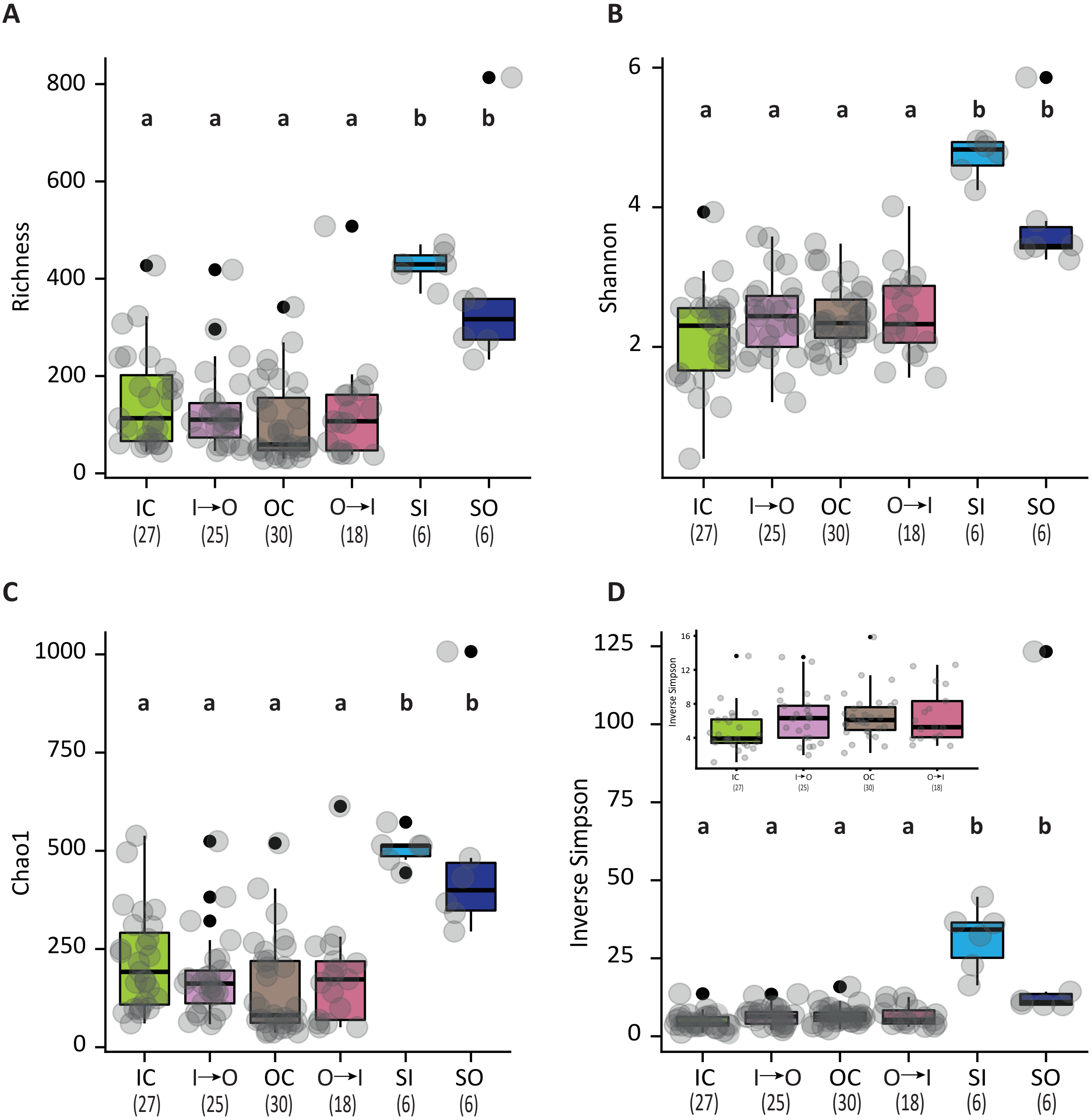

### Figure S2

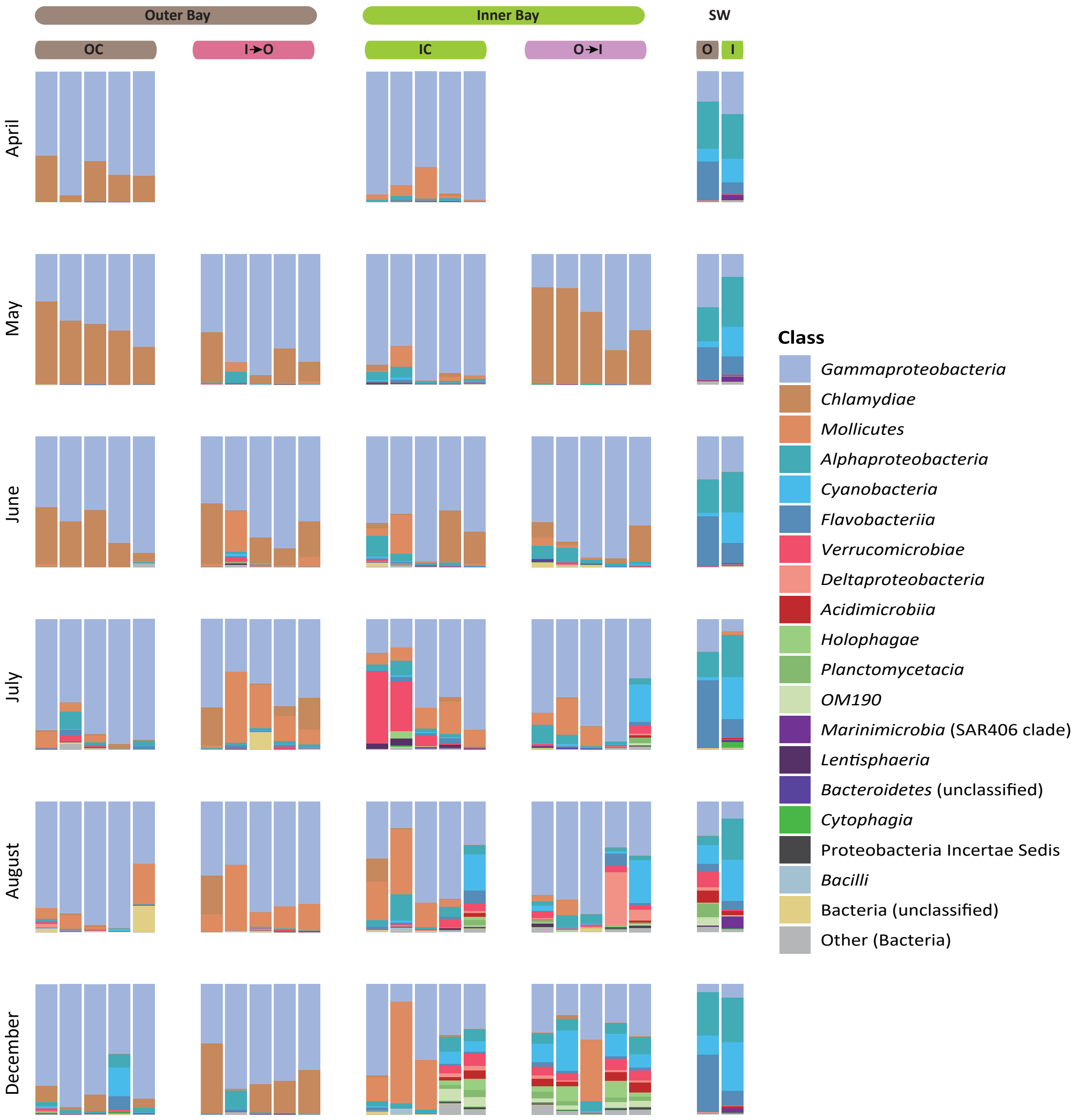

### Figure S3

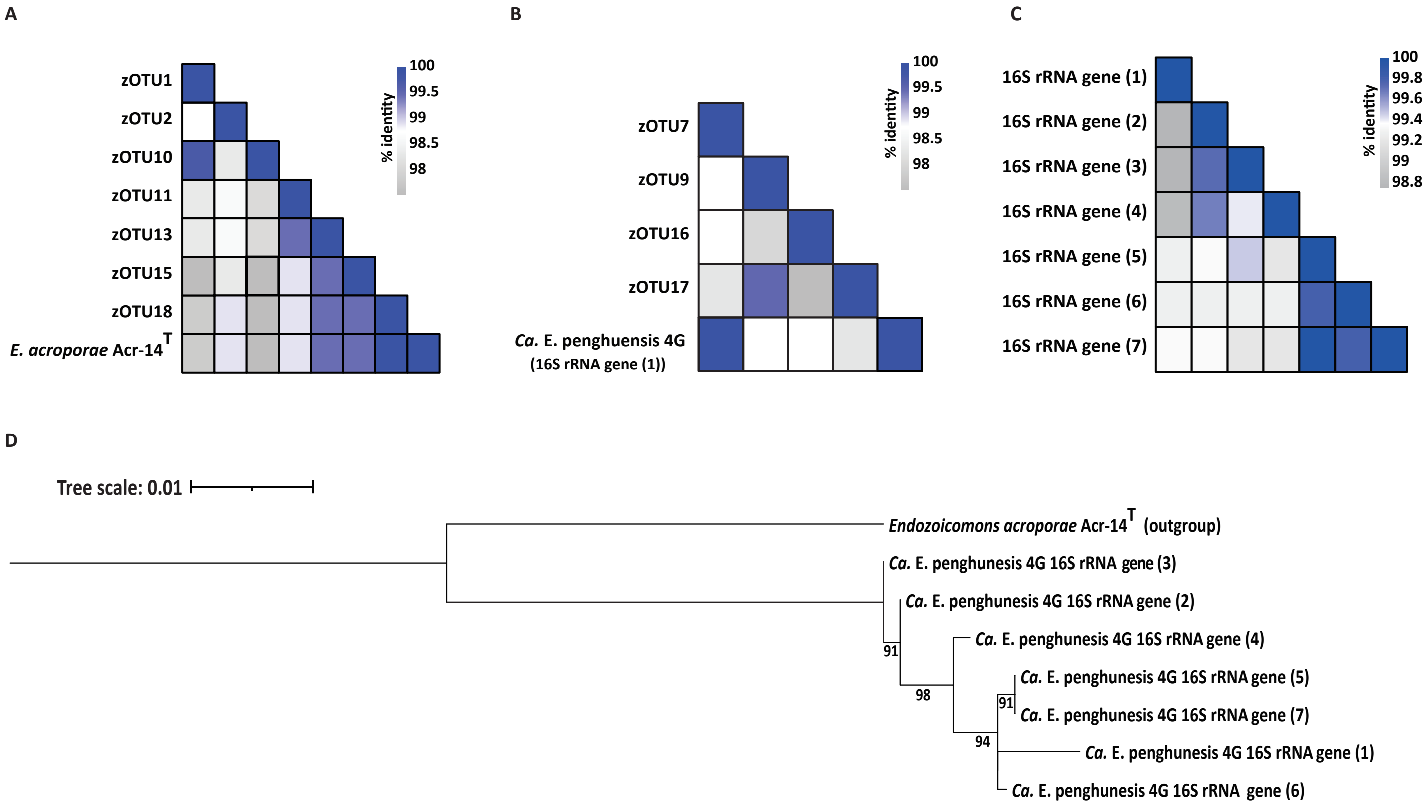

### Figure S4

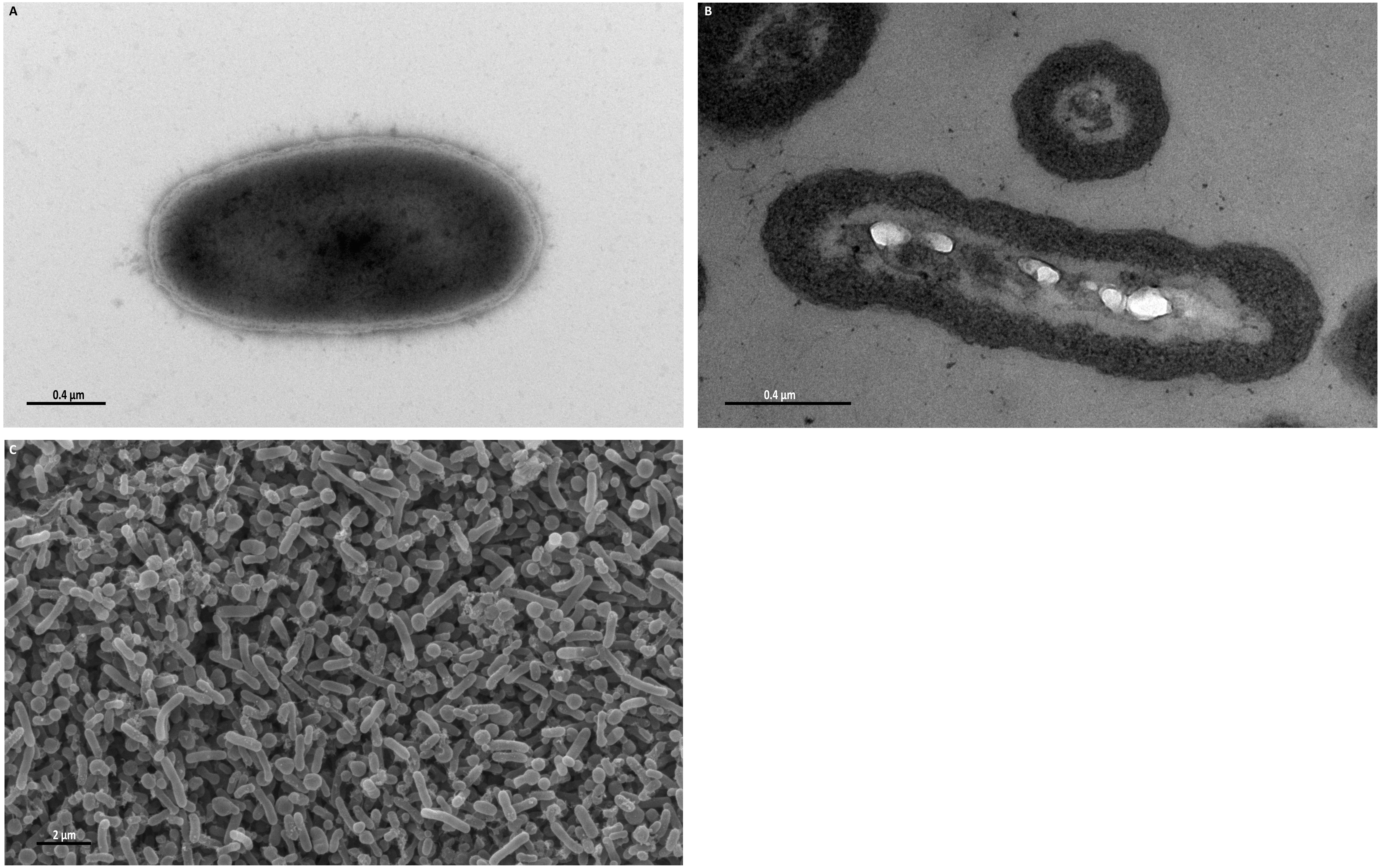

### Figure S5

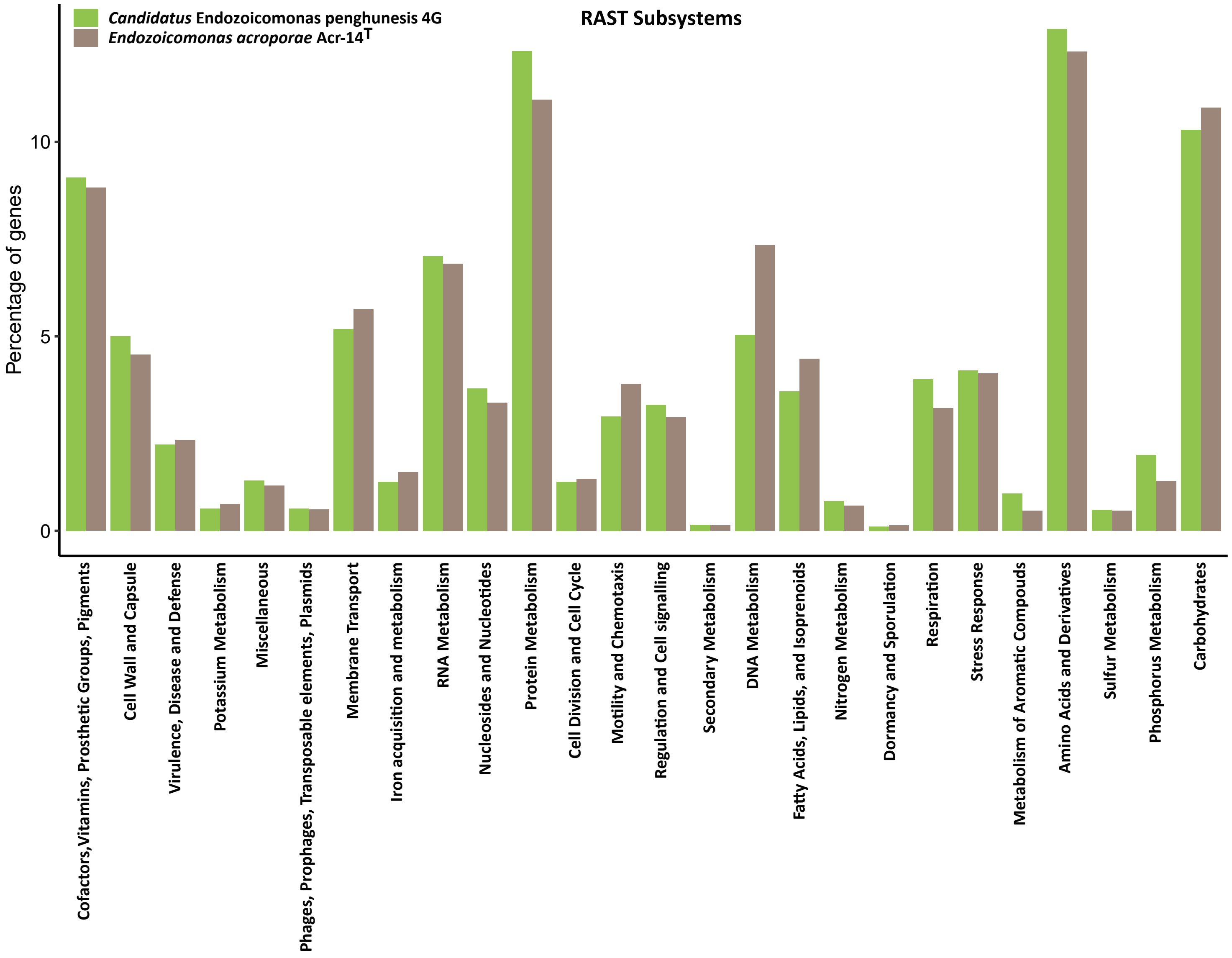

### Figure S6

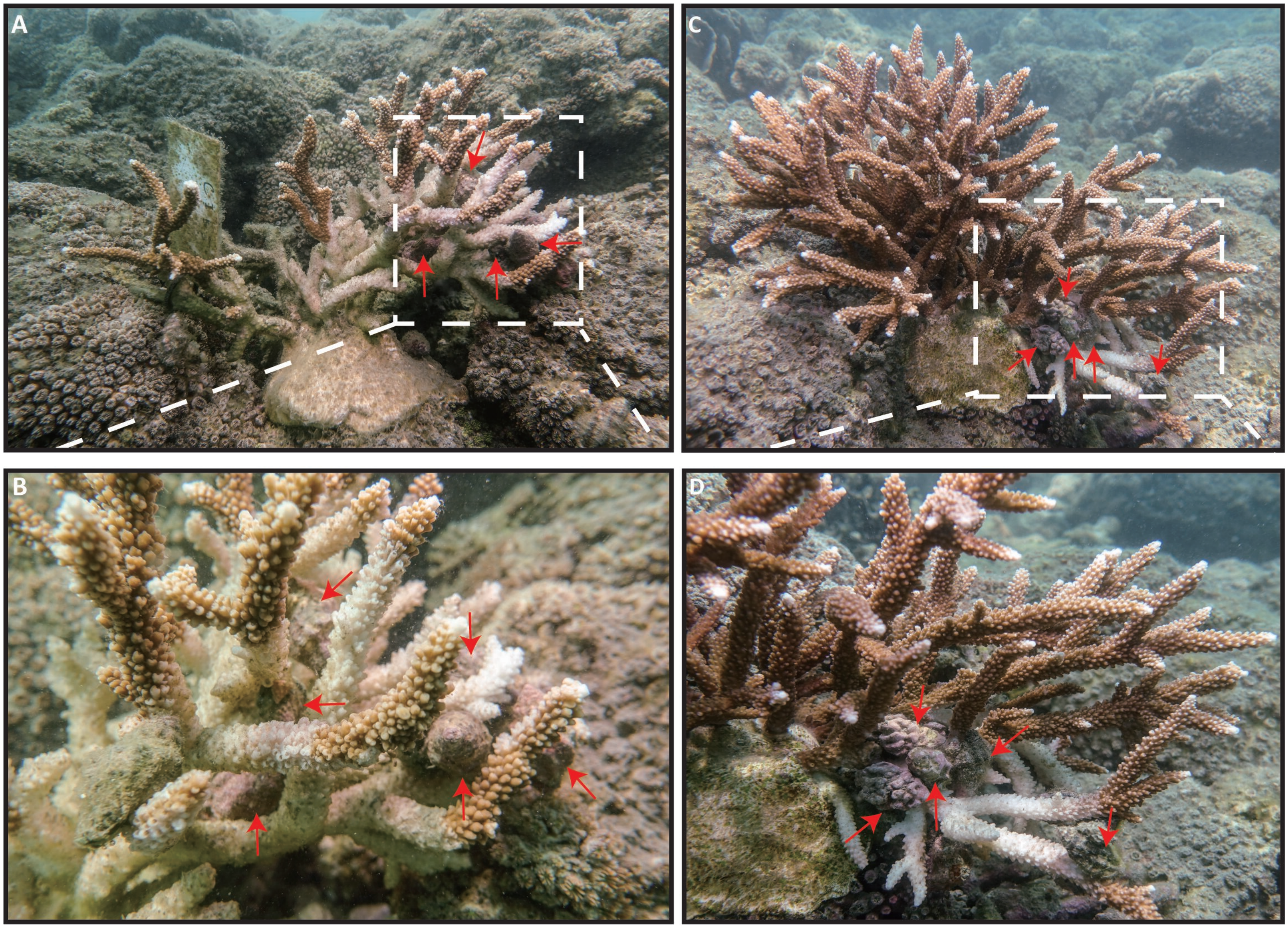

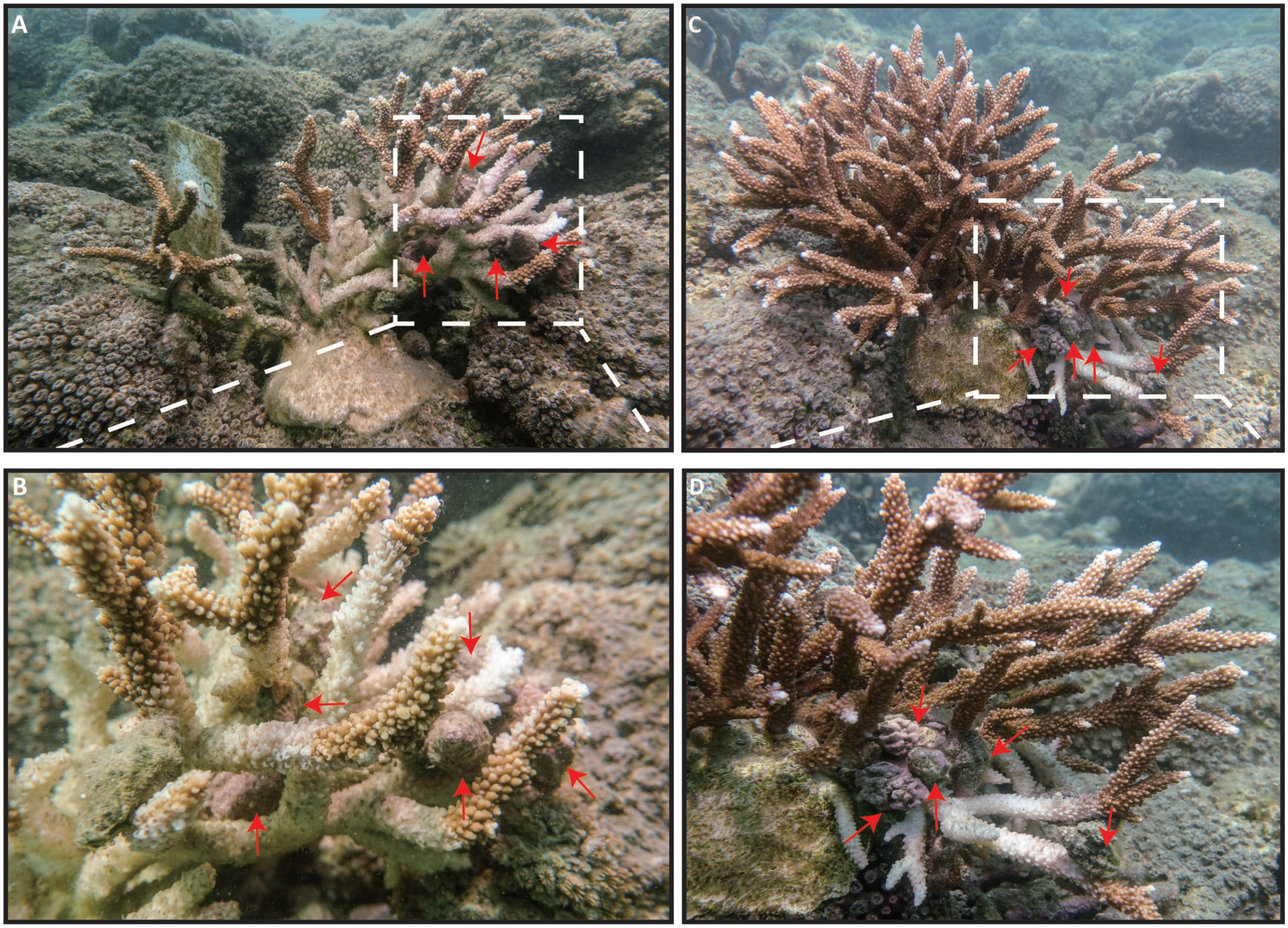
