## Supplementary material for "Microbiome restructuring: dominant coral bacterium *Endozoicomonas* species display differential adaptive capabilities to environmental changes": Table S1

Table S1. Recipe of modified marine broth (MMB)

| HEPES | 5.950 g/L |
| --- | --- |
| NaCl | 19.45 g/L |
| MgCl_2_$\cdot$6H_2_O | 18.79 g/L |
| Na_2_SO_4_ | 3.240 g/L |
| KCl | 0.550 g/L |
| CaCl_2_ | 0.120 g/L |
| NaHCO_3_ | 0.160 g/L |
| Peptone | 5.000 g/L |
| Yeast extract | 1.000 g/L |
| Trace element solution* | 1.000 ml/L |
| NP cocktail stock | 1.000 ml/L |
| pH | 7.20 |

***Trace element solution**

| FeCl_3_$\cdot$6H_2_O | 3.150 g/L |
| --- | --- |
| Na_2_EDTA | 4.360 g/L |
| CuSO_4_ primary stock | 1.000 ml/L |
| Na_2_MoO_4_ primary stock | 1.000 ml/L |
| ZnSO_4_ primary stock | 1.000 ml/L |
| CoCl_2_ primary stock | 1.000 ml/L |
| MnCl_2_ primary stock | 1.000 ml/L |

***Primary stock for preparing the trace element solution**

| CuSO_4_$\cdot$5H_2_O | 9.800 g/L |
| --- | --- |
| MnCl_2_$\cdot$4H_2_O | 180.0 g/L |
| Na_2_MoO_4_$\cdot$2H_2_O | 6.300 g/L |
| ZnSO_4_$\cdot$7H_2_O | 22.00 g/L |
| CoCl_2_$\cdot$6H_2_O | 10.0 0g/L |
