## Supplementary material for "Microbiome restructuring: dominant coral bacterium *Endozoicomonas* species display differential adaptive capabilities to environmental changes": Table S2

**Table S2. Differential phenotypic characteristics of *Candidatus* Endozoicomonas penghunesis 4G and *E. acroporae* Acr-14^T^.**

| **Characteristics** | ***Candidatus* Endozoicomonas penghunesis 4G** | ***Endozoicomonas acroporae* Acr-14^T *^** |
| --- | --- | --- |
| Colony pigment | Beige | Cream white |
| Size (µm) | 2.14 $\times$ 0.66 | 2.0 - 3.0$\times$0.5 -0.8 |
| Gram | Gram negative | Gram negative |
| Relation to O_2_ | FAN | Aerobic |
| Motility | $\pm$ | $-$ |
| Catalase activity | $-$ | $+$ |
| Oxidase activity | $+$ | $+$ |
| Enzymatic activities (API ZYM) |  |  |
| Alkaline phosphatase | $+$ | $+$ |
| Esterase | $+$ | $+$ |
| Lipase | $+$ | $+$ |
| Leucine arylamidase | $+$ | $+$ |
| Valine arylamidase | $+$ | $+$ |
| Cystine arylamidase | $+$ | $+$ |
| Trypsin | $+$ | $-$ |
| $\alpha$-chymotrypsin | $-$ | $-$ |
| Acid phosphatase | $+$ | $+$ |
| Naphtol-AS-BI- phosphophydrolase | $-$ | $+$ |
| $\alpha$-galactosidase | $-$ | $-$ |
| $\beta$-galactosidase | $-$ | $-$ |
| $\beta$-glucouronidase | $-$ | $-$ |
| $\alpha$ -glucouronidase | $-$ | $-$ |
| $\beta$-glucosidase | $-$ | $-$ |
| N-acetyl-$\beta$-glucosaminidase | $-$ | $-$ |
| $\alpha$-mannosidase | $-$ | $-$ |
| $\alpha$-fucosidase | $-$ | $-$ |
| **Sensitivity to antibiotic** |  |  |
| Streptomycin (10µg) | slightly sensitive | susceptible |
| Ampicillin (10µg) | susceptible | susceptible |

* (Sheu et al. 2017)

Sheu SY, Lin KR, Hsu MY, Sheu DS, Tang SL, Chen WM (2017) *Endozoicomonas acroporae* sp nov., isolated from *Acropora* coral. International Journal of Systematic and Evolutionary Microbiology 67:3791-3797
