## Supplementary material for "Microbiome restructuring: dominant coral bacterium *Endozoicomonas* species display differential adaptive capabilities to environmental changes": Table S3

**Table S3. *Candidatus* Endozoicomonas penghunesis 4G and related *Endozoicomonas* species strains growth condition**

| **Strain** | **Isolation location** | **Isolated source** | **Optimum/growth  temperature (**$\mathbf{℃}$**)** | **Optimum/growth salinity (PSU)** | **Optimum/growth pH** | **Reference** |
| --- | --- | --- | --- | --- | --- | --- |
| ***E. penghunesis* 4G** | Penghu, Taiwan | Hexacoral (*Acropora muricata*) | 20, 25 / 15 - 35 | 10, 20 / 5 - 30 | 8 / 6 - 9 | This study |
| ***E. acroporae* Acr-14^T^** | Kenting, Taiwan | Hexacoral (*Acropora* sp.) | 30 / 20 - 35 | 20 / 10 - 50 | 7 / 5 - 10 | (Sheu et al. 2017) |
| ***E. arenosclerae* Ab112^T^** | Rio de Janeiro, Brazil | Marine sponge (*Arenosclera brasiliensis*) | 20 - 30 / 12 - 35 | 30 / 20 - 50 | ND* | (Appolinario et al. 2016) |
| ***E. ascidiicola* AVMART05^T^** | Gullmarsfjord, Sweden | Sea squirt  (Scandinavian ascidians) | 23 - 25 / 5 - 27 | 10~20 / 5 - 50 | 6 - 7 / 6.2 - 8.3 | (Schreiber et al. 2016) |
| ***E. atrinae* WP70^T^** | Yeosu, Korea | Comb pen shell (*Atrina pectinata*) | 30 / 15 - 37 | 20 / 10 - 40 | 7 / 6 - 9 | (Hyun et al. 2014) |
| ***E. elysicola* DSM22380 ^T^** | Izu-Miyake, Japan | Sea slug (*Elysia ornata*) | 25 - 30 / 4 - 37 | >0 / >0 | ND | (Kurahashi & Yokota 2007) |
| ***E. euniceicola* EF212^T^** | Florida, USA | Octocorals  (*Eunicea fusca*) | 22 - 30 / 15 - 30 | 20 - 30 / 10 - 40 | 8 / 7 - 8 | (Pike et al. 2013) |
| ***E. gorgoniicola* PS125^T^** | Bimini, Bahama | Octocorals  (*Plexaura* sp) | 22 - 30/ 15 - 30 | 20 - 30 / 10 - 40 | 8 / 7 - 9 | (Pike et al. 2013) |
| ***E. montiporae* CL-33^T^** | Sourthern coast, Taiwan | Hexacoral  (*Montiporae aequituberculata*) | 25 / 15 - 35 | 20 - 30 / 10 - 30 | 8 / 6 - 10 | (Yang et al. 2010) |
| ***E. numazuensis* HC50^T^** | Namazu, Japan | Marine sponge | 25 / 15 - 37 | 20 / 10 - 50 | 7.5 - 8 / 5.5 - 9 | (Nishijima et al. 2013) |

*ND, not detected

Note: The optimum temperature of all strain are 20 - 30 $\mathbf{℃}$, the optimum salinity are 10 - 20 PSU and optimum pH are between 7 – 8.
